# Unbiased phosphoproteomics analysis unveils modulation of insulin signaling by extramitotic CDK1 kinase activity in human myotubes

**DOI:** 10.1101/2023.06.30.547176

**Authors:** Bitnara Han, Ginny X. Li, Wei Lin Liew, Edmund Chan, Shiqi Huang, Chin Meng Khoo, Melvin Khee-Shing Leow, Yung Seng Lee, Tianyun Zhao, Loo Chien Wang, Radoslaw Sobota, Hyungwon Choi, Mei Hui Liu, Kwang Pyo Kim, E Shyong Tai

## Abstract

Sensitivity and plasticity of insulin signaling and glucose uptake in skeletal muscle depends on determinants such as genetic variation and obesity. We collected muscle biopsies and isolated myoblasts from a multi-ethnic cohort of lean South Asians (N=10), lean Chinese (N=10), and obese Chinese (N=10), and analysed the proteome and phosphoproteome dynamics in terminally differentiated myotubes after a low-dose insulin stimulation (10nM at 0, 5, 30 min). The myotubes initially responded with increased abundance and phosphorylation level changes along the PI3K/AKT/mTOR axis, decreased abundance of translation initiation factors, and increased phosphorylation levels on proteins involved in mRNA processing at 5 min. After the acute response, protein abundance returned to baseline at 30 min, while phosphorylation changes persisted in proteins including AKT, RPS6 and AS160 (TBC1D4). A joint kinase-substrate statistical analysis revealed that protein abundance changes of AKT, PAK1 and CDK1 showed concordant phosphorylation changes in their respective substrates upon insulin stimulation. We also observed increased phosphorylation of some substrates uniquely in each group, particularly the substrates of CDKs showing stronger changes in South Asians than in Chinese. Pharmacological inhibition and siRNA knockdown of CDK1, a non-myogenic kinase, in terminally differentiated myotubes reduced glucose uptake and desensitized several phosphorylation-mediated signaling on protein translation initiation factors, IRS1, and AS160. Our data suggest that basal extramitotic activity of CDK1 is required for PI3K/AKT/mTORC1 signaling cascade and glucose uptake in insulin-stimulated myotubes. The data also provide a rich resource for studying the role of other kinases in the mechanism of insulin resistance in human myotubes.

## INTRODUCTION

Skeletal muscle is the main tissue responsible for insulin-stimulated glucose disposal ^1^. Ineffective cellular response to insulin in skeletal muscle cells is a key feature of insulin resistance (IR) and phenotypes within the metabolic syndrome and type 2 diabetes (T2D). IR is a complex phenotype with considerable variability between individuals, modulated in part by the ethnicity background during the onset with or without the interplay with obesity ^2–4^. Therefore, a proteome-scale phosphorylation map of insulin response dynamics in the skeletal muscle from multiple individuals provides a major data resource for identifying molecular components explaining the inter-individual variability in insulin sensitivity.

Past studies examining the impact of insulin stimulation on the global phosphoproteome have been acquired mostly in murine or human adipocytes and hepatocytes. The data from muscle fibers or differentiated myotubes are scarce likely due to the difficulty in clean isolation of myocytes from biopsies. The most popular workaround is to profile myoblasts^5, 6^. Batista and colleagues recently showed that skeletal myoblasts derived from induced pluripotent stem cells (iPSCs) of individuals with type 2 diabetes exhibited dysregulation of >1,000 phosphorylation sites when compared to myoblasts derived from individuals without type 2 diabetes, although most of these were insulin dependent ^6^. Hoffman and colleagues also described the global phosphoproteome changes via direct protein extraction from muscle biopsies after cycling exercise ^7^, yet the biopsies are expected to contain a mixture of multiple cell types and the experiments were performed on four healthy donors only.

In this study, we used quantitative mass spectrometry to investigate the phosphorylation-based insulin response in myotubes differentiated from thirty muscle biopsies. In particular, we aimed to study the variability of these changes across the individuals of different obesity status and ethnicity. Obesity is an established risk factor for T2D, strongly associated with the impairment of insulin-stimulated glucose uptake in skeletal muscle ^8^. Ethnicity may also play an important role in the development of IR. We have previously shown that obesity is inversely associated with insulin stimulated glucose uptake in general, but IR in South Asians was independent of obesity ^9^ whereas it was obesity dependent in Chinese individuals. South Asians exhibit IR even when they are lean. In contrast, Chinese are insulin sensitive when they are lean, but rapidly become IR with obesity, where the level of insulin sensitivity is indistinguishable from their South Asian counterparts at a BMI of 28 kg/m^2^ (which is not obese by the standards in population of European ancestry). We thus hypothesized that the global phosphoproteome profile of a sufficient number of myotube samples acquired from the three groups may reveal kinases and downstream signaling cascades underlying the variability among them.

Intriguingly, most published studies have reported that insulin stimulation triggers an array of downstream phosphorylation events not implicated in insulin signaling ^7, 10, 11^. For example, Humphreys and colleagues mapped phosphorylation events in 3T3-L1 adipocytes using global proteomics and phosphoproteomics and illustrated that a large proportion of the phosphorylated proteins was identified in substrates not related to insulin signaling pathway, suggesting that insulin regulates a number of other biological processes and cellular components ^11^. To understand this phenomenon better, we described our phosphoproteome data distinguishing the proteins that are annotated to be directly involved in the insulin signaling cascade from the ones that are not. Moreover, our data not only recapitulated the PI3K/AKT/mTORC1 signaling cascade, but also highlighted dynamic phosphorylation changes in the substrates of two other kinases, namely CDK1 and PAK1 upon insulin stimulation. Our follow-up experiment also shows that pharmacological inhibition of CDK1 phosphorylation activities and siRNA silencing of CDK1 directly result in reduced glucose uptake and desensitizes various components of the AKT signaling cascade in these terminally differentiated myotubes.

## RESULTS

### Primary human myoblast and myotube characterization

We harvested primary myoblasts from muscle biopsies of thirty individuals and differentiated them into myotubes (**Figure 1a, STAR Methods**). The participants consist of three distinct groups of ten individuals: 10 obese Chinese (CO), 10 lean Chinese (CL), and 10 South Asian lean participants (SL). We chose myotubes instead of the initially harvested myoblasts because they were more sensitive to insulin stimulation as measured by AKT phosphorylation, especially at low doses of insulin (**Supplementary Figure 1a**). Immunoblotting of phospho-AKT (serine residue 473) on all 30 myotubes exhibited time-dependent increases in response to insulin **(Figure 1b)**, indicating successful acute stimulation.

**Figure 1.**
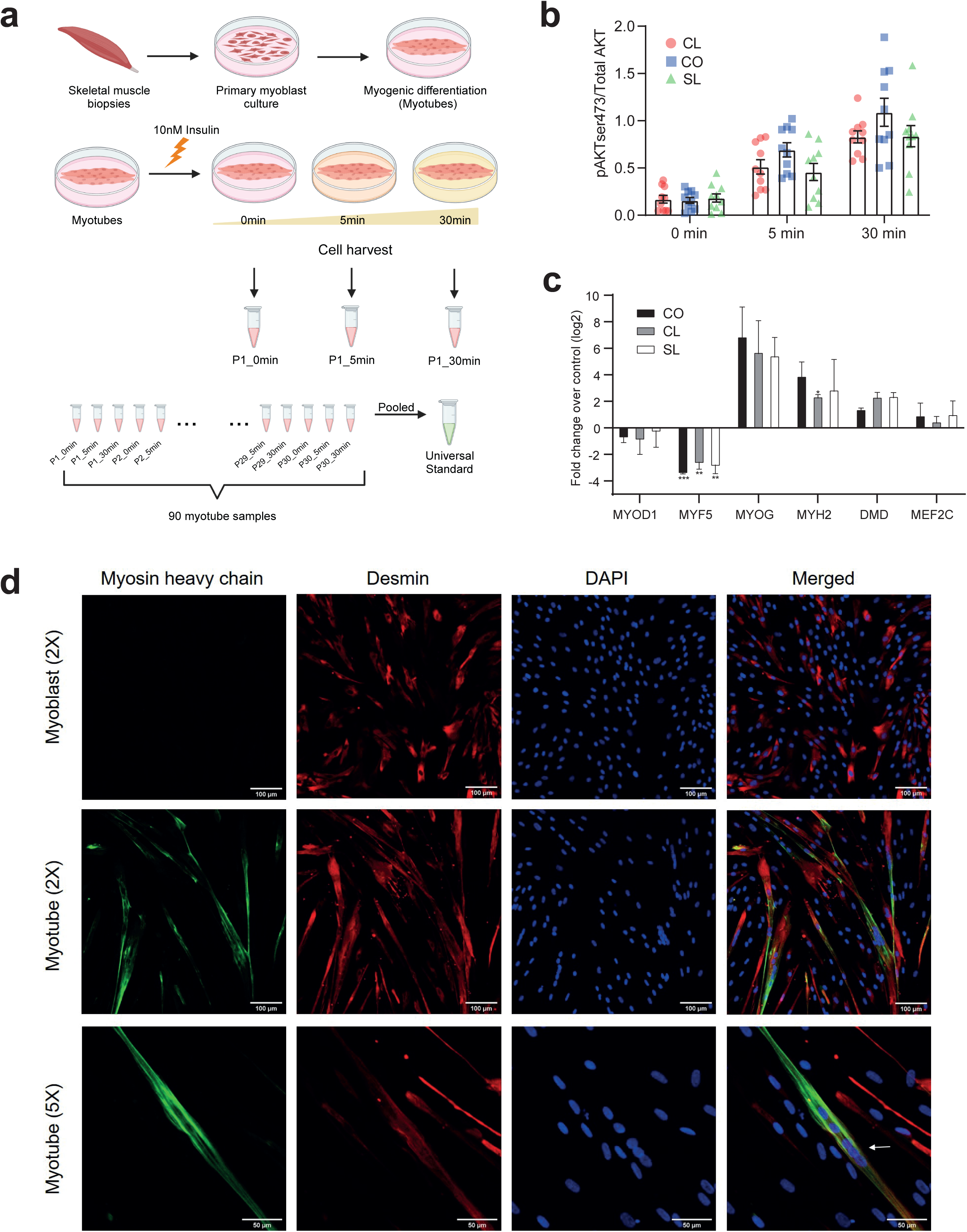
**(a)** Myotube cell lines extracted from muscle cells of all 30 patients were treated with 10 nM insulin, respectively, and cells were harvested at 0 minutes and 5 minutes and 30 minutes. Universal standard was used to correct the variation between experimental batches, which was made by pooling all of the proteins extracted from all 90 samples. **(b)** Densitometric analysis of p-AKT protein levels in human myotubes isolated from CL, CO and SL subjects (n=30, 10 from each group) at basal level (0 min) or after stimulation (10 nM insulin) at 5 and 30 minutes. Graph displays mean p-AKT levels in arbitrary units (AU) normalized against total AKT protein levels. Error bars are expressed as SEM. **(c)** Log2 fold change of early, middle and late stage myogenesis markers in day 7 myotubes compared to day 0 myoblasts control. Results were normalized to *RPLP0* expression level and expressed as mean ± SEM (n=9). *:p<0.05, **p:<0.01 and ***p:<0.001 with paired Student’s t-test. **(d)** Immunostaining of representative myoblast and myotube cultures at different magnifications (2X or 5X digital zoom on Olympus Fluoview software). Desmin (red) and fast skeletal myosin heavy chain (green) was stained with Alexa 555 and 488 respectively, counterstained with DAPI (blue). White arrow shows myotube with multiple fused nuclei.

Gene expression and immunofluorescence analyses were also performed to characterize a subset of the myotubes from each group. Compared to the myoblast stage, the myotubes showed decreased gene expression of early-stage myogenic differentiation markers, MYOD1 and MYF5 and increased expression of middle to late-stage markers, such as MYOG, MYH2, DMD and MEF2C (**Figure 1c**). Immunofluorescence staining of cytoplasmic desmin in the myotubes indicated an elongated and linear cell morphology. The myotubes were also multi-nucleated, an indication of myogenic cell fusion, and expressed fast myosin heavy chain, a marker for myogenic differentiation (**Figure 1d**). In all, we showed that these myotubes were between middle to late stage of myogenic differentiation and have a cell morphology that was distinct from the myoblast stage. Lastly, we show that insulin stimulated glucose uptake responses of the myotubes were between 1.2 to 2.4 folds across groups (**Supplementary Figure 1b**), consistent with previous reported ranges ^12^.

### Global landscape of protein abundance and phosphorylation signaling dynamics

We next treated the myotubes with 10 nM of insulin and sampled cells at the time of stimulation (0 min) and 5 and 30 min post stimulation. We performed isobaric tandem mass tag (TMT) labelling-based quantitative proteomics analysis of these cells to identify insulin-dependent signaling pathways that may be differentially modulated across these groups (**Supplementary Figure 2**). We deliberately treated the myotubes with a low dose of insulin in this experiment, which we deemed a level that not only stimulated the cells sufficiently but also captured subtle differences across the three groups that might otherwise be masked by effects of saturated insulin signaling (**Supplementary Figure 1b**).

The global proteomics experiment identified a total of 8,964 proteins across the 16-plex TMT experiments. Among these, 8,013 proteins (89%) were commonly detected and quantified across all TMT sets. Meanwhile, the phosphoproteomics experiment identified 11,715 phosphorylation sites on 3,293 proteins, reporting 3.56 phosphorylation sites per protein on average. We have also benchmarked our proteome and phosphoproteome coverage against published data sets, and we have detected a comparable number of proteins and phosphorylation sites to the previous data sets (see the comparisons in **Supplementary Information**).

In response to stimulation, protein abundance increased from the basal levels in approximately 14% and decreased in about 19% of all detected proteins (1,157 and 1,555 out of 8,013) at 5 min (local false discovery rate ≤ 0.05). Proteins directly involved in the PI3K/AKT signaling cascade showed increased abundance in general (**Figure 2a**), including the catalytic subunit of phosphatidylinositol 3-kinase PIK3CA, isoforms of protein kinase B (AKT), mTOR regulatory protein RPTOR (also known as raptor for mTORC1). Overall, the protein abundance changes were modest across the subjects, with fewer than 12% of the proteins showing absolute fold changes greater than 10%. The most statistically significant changes at 5 min include decreased abundances of Ras oncogene homolog proteins (HRAS, KRAS, NRAS), integrins (ITGA1, ITGA3, ITGAV, ITGB3, ITGB5), and 14-3-3 chaperones (YWHAB, YWHAE, YWHAG, YWHAH, YWHAQ, YWHAZ). These proteins have not been described to be first line responders in insulin signaling nor do they carry out directly related biological functions. At 30 min, protein abundance mostly recovered to the baseline levels (**Figure 2b**). Key proteins in the insulin signaling pathway did not show abundance changes compared to the control state at 30 min. However, among the remaining proteins with average fold changes below 10%, one in five proteins showed absolute fold changes greater than 10% in at least one of the subgroups (CO, CL, SL), indicating group-specific protein abundance changes (**Supplementary Table 2**).

**Figure 2.**
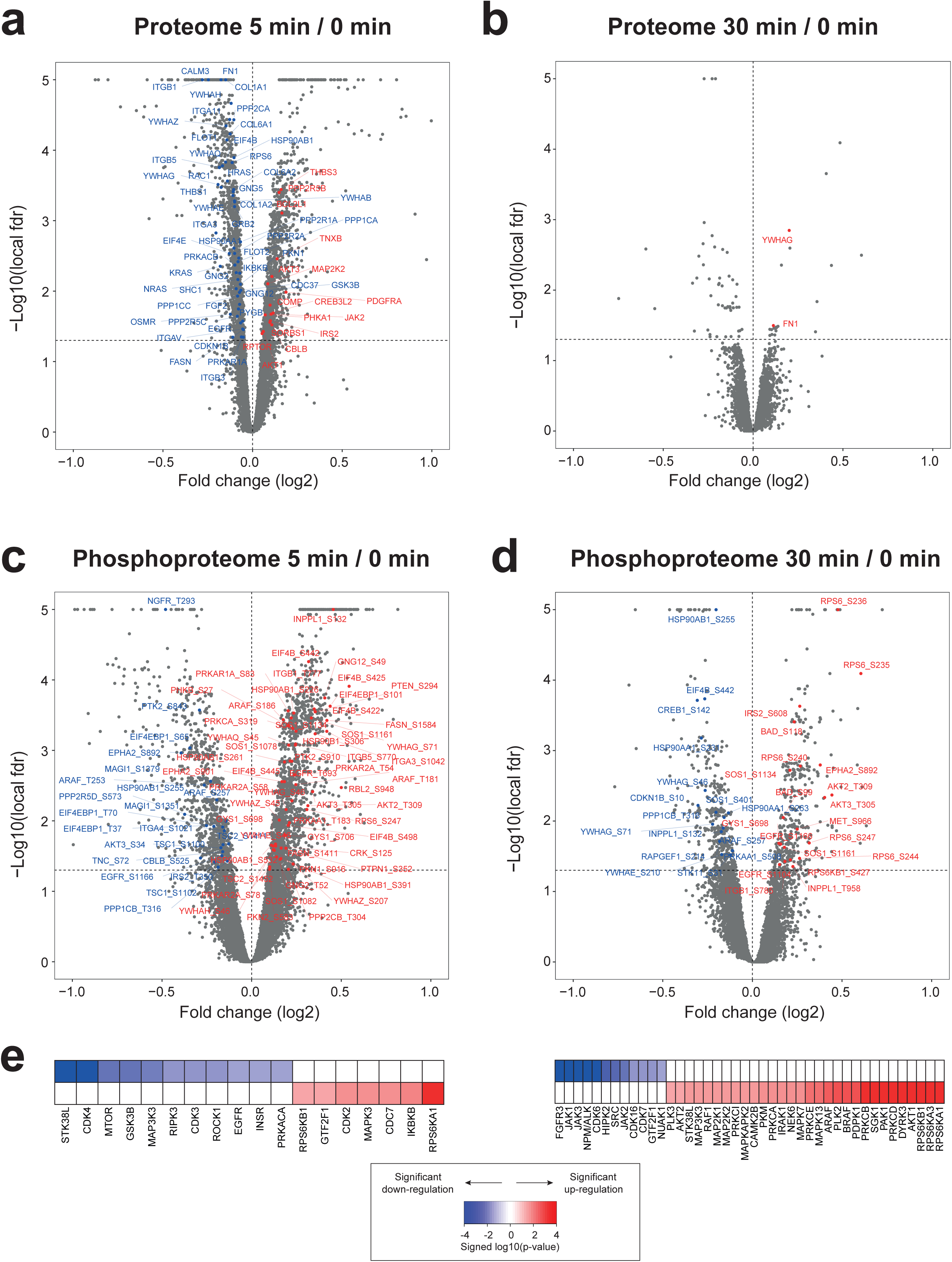
**(a-d)** Volcano plots of log2 fold change and -log10 P value (cutoff > 2, or p-value 0.01) at the two time points in the proteomics and the phosphoproteomics data. Red or blue dots are proteins or phosphorylation sites with statistically significant changes (local false discovery rate < 0.01) in the insulin signaling pathway and the PI3K/AKT signaling pathway: red and blue are increased or decreased abundance or phosphorylation, respectively. (e) Kinases enriched by concordantly increased or decreased phosphorylation levels at 5 min and 30 min, reported by KSEA.

Meanwhile, the phosphoproteome showed changes of greater magnitude to the insulin stimulation than the proteome at both time points (**Supplementary Table 3**). At 5 min, we observed significant increase in the phosphorylation levels in about 15% and decrease in 8% of all detected phosphorylation sites (1,743 and 939 out of 11,715) (**Figures 2c and 2d**). These include expected phosphorylation changes including increased phosphorylation levels in AKT2, the AKT isoform most abundant in skeletal muscle,^13^ the GDP/GTP exchange promoter SOS, the translation initiation factor EIF4B, and the chaperone protein HSP90AB1. Decreased phosphorylation was most prominent in certain residues of tuberous sclerosis complex protein TSC1 and membrane-associated guanylate kinase MAGI1 as well as sites on insulin receptor substrate 2 (IRS2), all of which have implications for their context-specific roles via activation or inhibition of their respective targets. Interestingly, many proteins showed both increased phosphorylation and decreased phosphorylation at multiple positions, including translational initiation factor 4E binding protein (EIF4EBP1), and epidermal growth factor receptor EGFR. This observation suggests that efficient phosphorylation-mediated signaling on individual agents of the whole cascade likely requires coordination of multiple kinases at adjacent target residues through their collective impact on the substrate protein conformation.

At 30 min, 3% and 4% of the phosphoproteome exhibited increased and decreased levels, respectively (391 and 464 of 11,715). Although we observed withering responses across multiple phosphorylation sites over time, many sites maintained elevated phosphorylation levels, especially on the proteins directly involved in the PI3K/AKT signaling cascade including AKT (AKT2/AKT3) and IRS2, as well as others that are further downstream of the direct signaling cascade such as SOS and RPS6. We unfortunately did not detect any phosphopeptide in AKT1 isoform in our experiment likely due to the low abundance in myotubes, but we confirmed serine 473 phosphorylation on AKT1 or serine 474 on AKT2 for all samples by western blot (see **STAR Methods**).

In summary, our data show that the insulin stimulation induced acute protein abundance changes of moderate magnitude at 5 min and restored the pre-stimulation levels at 30 min. Phosphorylation response was detectable immediately at 5 min with levels increased and decreased phosphorylation in and out of the insulin signaling pathway. Some of these responses persisted through 30 min, which may be the result of the increased abundance and the heightened activity of certain kinases at 5 min. In mammalian cells, the presence of early and late phases in response to growth factor cues has been increasingly noted in global proteomics data. Golan-Lavi et al. have previously shown the diversity in transient increase and decrease of protein abundance in human mammary cells in response to an epidermal growth factor ^14^. Collins et al. have also demonstrated that the perturbation of the interactome of 14-3-3β scaffold protein by IGF1 treatment was manifested in different groups of proteins with early and late changes, with early changes occurring within 1 min ^15^.

### Substrate phosphorylation changes encompass pathways beyond insulin signaling

We next examined the biological pathways significantly enriched in the proteome and phosphoproteome changes at both 5 min and 30 min (**STAR Methods**). At 5 min, we observed increased protein abundance in chromatin modifying enzymes such as histone acetylases (HATs) and deacetylases (HDACs), elongated acetyltransferase complex (ELPs), lysine demethylases (KDMs) and ATP-dependent transcription activators (SMARCs). In addition, proteins involved in P53 degradation (AKT1, AKT3, ATM, CDK1, CDKN2A, DAXX and RICTOR) and processes governing inositol phosphates and its intermediates (IMPA2, INPP4A, INPP4B, PLCB1, PLCD1, PLCD4) consistently increased in abundance (**Supplementary Table 4**).

Meanwhile, the proteins with decreased abundance at 5 min also showed enrichment of a broad spectrum of biological processes including gene expression and protein metabolism, indicating that the cells responded to the insulin stimulation by halting cell proliferation during the initial phase of the response (**Supplementary Table 4**). This includes members of the 14-3-3 family, core phosphoprotein-binding chaperones that mediate glucose uptake ^16^. These chaperones are part of other pathways enriched for the proteins with reduced abundance at 5 min, including insulin-mediated glucose transport, translocation of GLUT4 to the plasma membrane, vesicle-mediated transport and membrane trafficking, and apoptosis (**Supplementary Table 4**).

To interpret the phosphoproteome dynamics in connection with kinase activity, we conducted a kinase-substrate enrichment analysis (KSEA) (**Supplementary Table 5**). The substrates of p90 ribosomal S6 kinase subunits (RPS6KB1, RPS6KA1), inhibitor of nuclear factor IKBKB, MAPK3, CDK2, cell division cycle kinase CDC7 and general transcription factor GTF2F1 show increased phosphorylation at 5 min, whereas the substrates of CDK3, CDK4, mTOR, protein kinase C PRKCA, and ROCK1 showed decreased phosphorylation levels at 5 min (**Figure 2e**). At 30 min, we see significant increase of phosphorylation in the substrates of insulin-related kinases such as AKT1, AKT2, RAF1, PAK1, BRAF, MAPK7, and MAPK13, suggesting activated canonical insulin signaling at 30 min. This timing also aligns well with the priming time reported in the adipocytes in Humphreys et al. (20 min after 100 nM treatment).

### Kinase-substrate modelling captures PAK, MAPK, and CDK as central kinases in the extended signaling network

It is well known that substrate phosphorylation site occupancy and subsequently the overall kinase activity are not solely determined by the kinase protein abundance but by a multitude of other factors such as cellular localization of kinases and substrates as well as detailed conformational changes of substrate phosphorylation sites ^17, 18^. It is nevertheless plausible that kinase abundance indeed affects the activity in certain human kinases^19^, and this relationship has not been thoroughly investigated in specific cell types such as myotubes in dynamic conditions. For this reason, we next focused on the kinase protein abundance changes at 5 min with concordant phosphorylation changes at 5 min and 30 min to identify concurrently responsive kinase-substrate pairs, assuming that the abundance changes in the kinase proteins correlated with their corresponding substrate phosphorylation levels are more specific responses to insulin stimulation than the substrate phosphorylation level changes without the concurrent change in the kinase. To this end, we implemented a novel Empirical Bayes method for 2D (or paired) differential analysis of kinase-substrate, called KSA2D (**STAR Methods**), which identifies concordant changes between kinase abundance and substrate phosphorylation in the same samples.

We collected a map of 16,718 high confidence kinase-substrate phosphorylation events from multiple data sources and applied KSA2D to all 30 samples (**STAR Methods**). We selected 658 and 415 kinase-substrate pairs (**Supplementary Table 6**) as jointly differential pairs respectively (probability≥0.95, estimated local false discovery rate 0.05). 237 of the 658 pairs (36%) showed concordant increase in kinase abundances at 5 min and substrate phosphorylation levels at 5 min, and 150 of the 415 pairs (36%) showed concordant increase in kinase abundances at 5 min and substrate phosphorylation levels at 30 min. The substrate phosphorylation sites identified by this analysis clearly demonstrated enrichment of the minimal kinase motifs containing kinase specificity-determining residues proline at +1 position and arginine at -3 position (**Supplementary Figure 3**) ^20^.

Investigating the concordant kinase abundance changes at 5 min and substrate phosphorylation levels at 5 min, KSA2D analysis of all thirty subjects identified kinases known to be directly involved in insulin signaling: AKT, MTOR, RPS6KB1, PRKACA, CDK4, MAPK3 and MAP2K2 (orange edges in **Figure 3a**). It also highlighted cyclin-dependent kinase CDK1 as the most active kinase in which the protein abundance increase was correlated with the elevation in the phosphorylation levels of substrates. However, many of those substrates were not the best characterized substrates of the CDK1/cyclin B complex during cell cycle, suggesting that extramitotic CDK1 activity can be invoked upon insulin stimulation in myotubes. These substrates were mostly involved in non-insulin signaling functions, including protein synthesis and cell division. Interestingly, CDK1 ^21^ is known to phosphorylate serine residues on RPTOR ^22^ and p70S6 kinase RPS6KB1 ^23^, indicating that the kinase may have an intrinsic modulatory role through regulation of mTORC1 activity during insulin signaling in myotubes, which we revisit later in the Results section.

**Figure 3.**
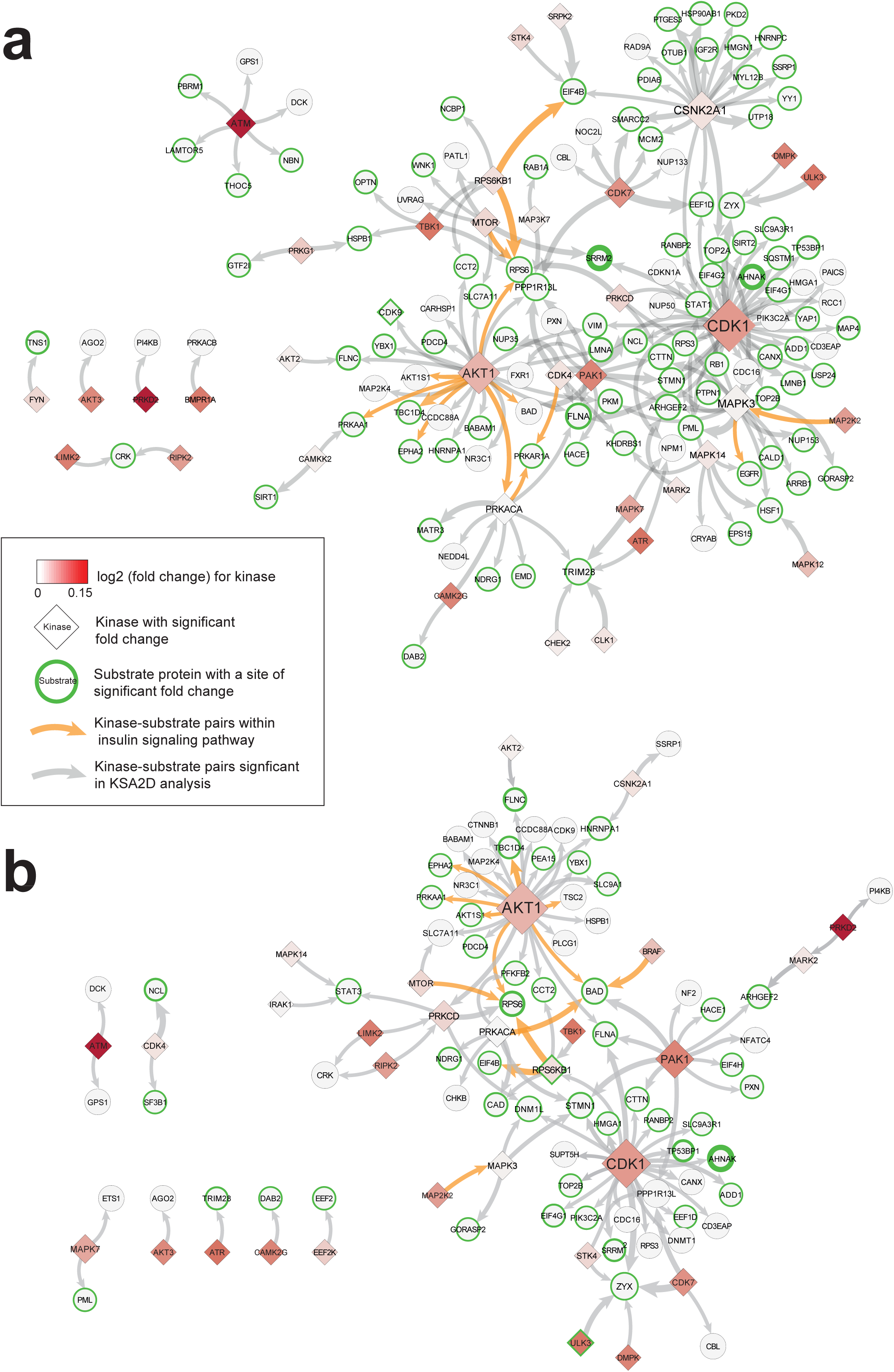
Network of kinase-substrate pairs with significant positive co-variation in all thirty subjects (2D analysis results, at local false discovery rate 0.05). Significant relationships of 5 min kinase with 5 min substrate in **(a)** and 5 min kinase with 30 min substrate in **(b).** Each edge represents phosphorylation of a substrate (sites omitted, in round circles) by a previously validated kinase (in diamonds). Edge thickness is proportional to the number of possible phosphorylation sites seen in our data between the kinase and the substrate protein. Node color in diamonds shows kinase protein abundance change. Green label represents significant kinase abundance change. Green node border indicates occurrence of significant site level change of the corresponding substrate, and the border thickness is proportional to the number of significant sites. Label and node size are proportional to the degree of connectivity in the network. Orange edge represents known kinase-substrate relationship in insulin signaling pathway.

In addition, PAK1, the serine/threonine-protein kinase, was also found to be a hub kinase potentially phosphorylating multiple residues on substrates including CTTN, PPPIR13L and BAD. BAD was the common target of both AKT1 and PAK1. Murine PAK1 has previously been implicated as a required element in GLUT4 recruitment in mouse skeletal muscle *in vivo* ^24, 25^. Tunduguru *et al*. ^26^ have also found that PAK1 signaling promotes insulin-stimulated GLUT4 vesicle translocation to the cell surface by contributing to cortical actin remodeling ^27^.

We next investigated the association between the kinase abundance changes at 5 min and the substrate phosphorylation level changes at 30 min (**Figure 3b**). We again observed a high correlation between the abundance changes of the kinases involved in insulin signaling (AKT, MTOR, RPS6KB1, PRKACA, MAP2K2 and BRAF) and increased phosphorylation of their known substrates such as TBC1D4, RPS6, EPHA2 and EIF4B. Phosphorylation on BAD (S99, S118) increased significantly at 30 min, which is a common substrate of AKT1, PRKACA, PAK1 and BRAF. CDK1 and PAK1 remained to be hub kinases, likely responsible for the phosphorylation of filamin A (FLNA) and cortactin (CTTN).

We refer the readers to **Supplementary Figure 4** for a more detailed view of the signaling events identified through KSA2D in the context of specific biological processes (green edges represent concordant increase in both kinase abundance and phosphorylation levels).

### Variability of phosphorylation changes across the weight and ethnicity groups

After exploring the response shared by all thirty individuals, we next examined the patterns in the kinase protein abundance and substrate phosphorylation changes in each of the three groups. Consistent with the analysis of all 30 samples, more concordant activities were observed at 5 min than at 30 min in all the subgroups. However, KSA2D analysis reported a number of group specific changes in the kinase abundance and substrate phosphorylation (**Supplementary Figure 5**). For example, the phosphorylation changes in the AKT isoforms and PAK1 substrates were more pronounced in Chinese obese group (CO), whereas the phosphorylation on the substrates of CDK1 were more pronounced in South Asian lean group (SL). The kinase abundances of ERKs (MAPK3, MAPK1) increased at 5 min while the phosphorylation levels of their common substrate EGFR increased at 5 min simultaneously in SL only. The kinase abundance levels of BRAF, AF1, PAK1, AKT and PRKACA increased with their shared substrate phosphorylation level of the residues on BAD at 30 min.

We next compared the fold change data (logarithmic scale, base 2) for protein abundance and phosphorylation globally between all pairs of groups. Due to the limited sample size (N=10 per group) and mild effect sizes, we did not have the statistical power to detect many statistically significant differences between groups. As such, we instead examined obesity-associated differences in the fold changes between CO and CL, and ethnicity-associated differences in the fold changes between SL and CL (log2 fold change differences of 0.2 or greater) (**Supplementary Figure 6**, red text indicating log2 fold changes > 0.2; **Supplementary Table 7**). The group-to-group differences were considerably more pronounced in the phosphorylation data than in the protein abundance data at both time points. For example, the phosphorylation levels of residues on IRS2, CBLB, EIF4EBP1 and PRKAR2A at 30 min remained mildly decreased from baseline in CL, but increased in CO, hinting at different plasticity of upstream regulators of the signal transduction between the two groups. The phosphorylation levels of multiple serine residues on ERBB2, the receptor engaging with IRS proteins and thus reducing their interaction with the insulin receptor ^28^, decreased at 30 min in CO but increased in CL. A similar contrast was also observed on BAD, known to be phosphorylated by PAK1, PAK4 and PRKACA and inhibit apoptosis induction thereafter ^29^, suggesting that these proteins may confer differential sensitivity to a mild dose of insulin stimulation between obese and lean individuals.

Interestingly, we observed enrichment of Rho GTPase-activating ROCK/PAK pathways in most proteins with increased phosphorylation at 30 min uniquely in CO, including MYH9, PPP1R12A, ROCK1 and FLNA. By contrast, the phosphorylation levels of proteins in cell adhesion-associated pathways remained stagnant at 30 min in CO but increased in CL. These genes include AFDN, CTNND1, and CDH13, all implicated in adherens junction interactions, which directly regulates actin cytoskeleton. In addition, phosphorylation of CTNND1, CTTN, MET and PLCG1 also increased at 30 min in CL but not CO, all involved in fibroblast growth factor (FGF) signaling pathway. FGF is an essential growth factor for attenuating insulin resistance ^30^, and aberrant FGF signaling has been implicated in skeletal diseases ^31^, reflecting a differential plasticity to mobilize other growth factor signaling pathway in the lean subjects.

The global data also showed phosphorylation changes uniquely observed in South Asians but not in Chinese subjects. Phosphorylation levels on the proteins involved in the apoptotic execution phase (ADD1, DBNL, PAK2, PTK2, TJP2, VIM) increased from baseline in SL and decreased in CL at 5 min. By contrast, the phosphorylation levels of proteins in other pathways decreased in SL, but increased in CL at 5 min. These include interleukin-4 signaling pathway (BAD, IRS2, FLNA, SOS1 and STAT5A) and cell junction organization (AFDN, CDH15, CTNND1, CTNNA1, FLNA, FLNC, PLEC, PXN). At 30 min, we observed increased phosphorylation of PAK2, CFL1, NRP1 in SL, all involved in PAK-dependent axon repulsion, and decreased phosphorylation on the same sites in CL. A similar contrast was observed in the proteins involved in Rho GTPase-activating PAK pathway, including PAK2, CTNN and NF2.

### Cdk1 inhibition desensitizes glucose uptake and phosphorylation levels of AKT and mTOR substrates

Given the extensive phosphorylation changes in the CDK1 substrates outside cell division regulation, we next set out to characterize the role of non-mitotic kinase activity of CDK1 in insulin signaling through pharmacological inhibition and translational repression via siRNA-based knockdown. Specifically, cultured myotubes were treated with RO-3306, a selective ATP competitive inhibitor to inhibit CDK1 activity, or transfected with a CDK1 targeting siRNA pool to decrease CDK1 expression. Glucose uptake is one of the primary outcomes of insulin signaling and is especially pertinent to muscle as it is the main tissue responsible for glucose disposal. Inhibition of CDK1 resulted in a decrease in insulin stimulated glucose uptake compared to the respective control, while the same effect was similarly observed in the CDK1 knockdown samples, albeit more mildly **(Figure 4a)**. This observation further corroborates the importance of CDK1 in the signaling cascades governing glucose uptake under insulin stimulated conditions. To further delineate how altered CDK1 activity and expression can impact the insulin signaling pathway, we treated the CDK1 inhibited and knockdown samples and their respective controls with the same low dose of insulin (10nM) with time-course sampling at 0, 5 and 30 min post stimulation. We then carried out quantitative phosphoproteomics analysis to identify differences in the phosphorylation network between the treated and control samples (**Figure 4b**). Due to the limited sample sizes in this experiment compared to the main data (N=3), we used a well-established empirical Bayes method (EBprot) that does not depend on hypothesis testing to select significantly differentially altered phosphorylation sites (FDR 5%) ^32, 33^.

**Figure 4.**
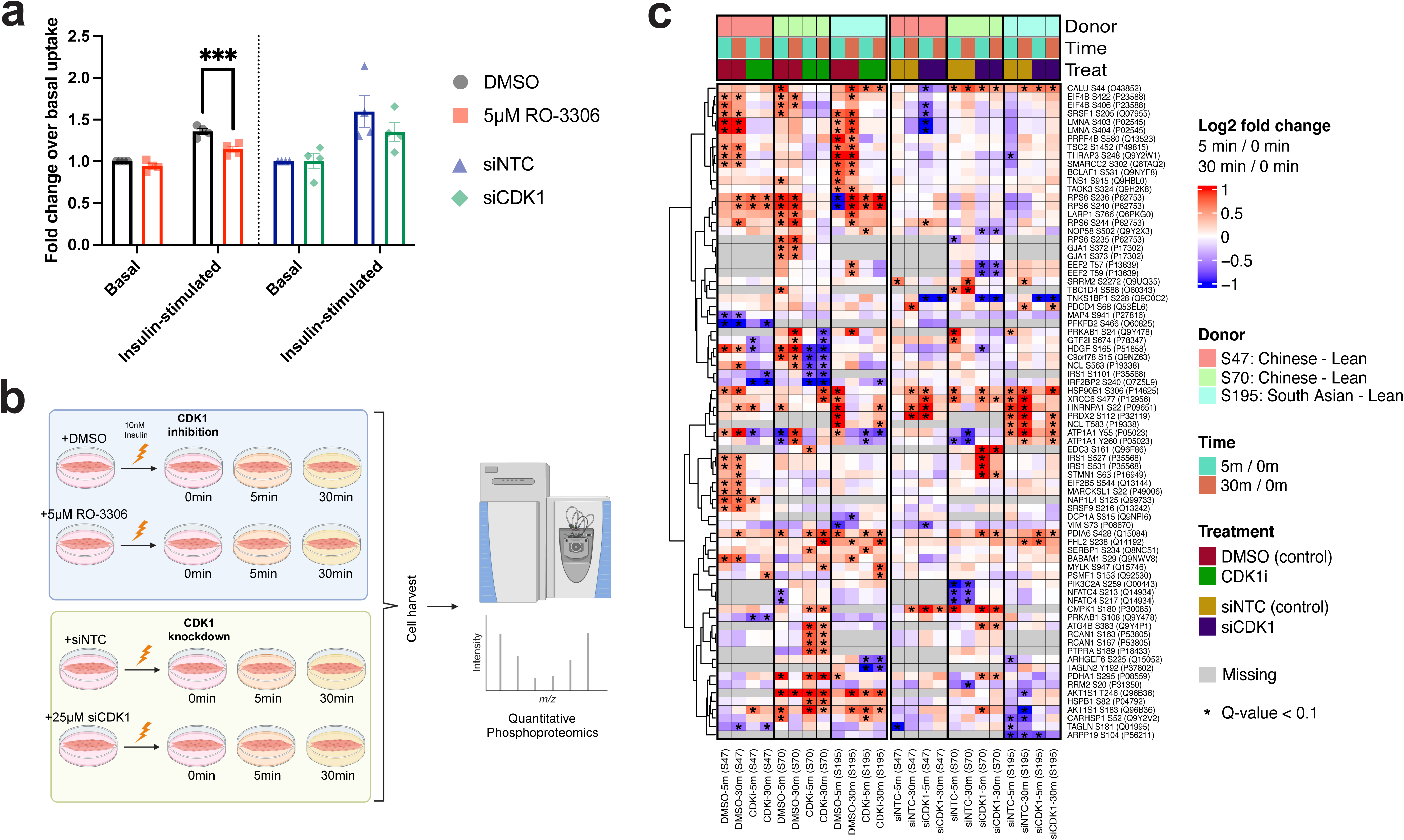
**(a)** Insulin-stimulated glucose uptake quantified by measuring luminescence signal proportional to concentration of 2DG taken up by myotubes. Data is presented as fold change over respective basal uptakes and expressed as mean ± SEM (n=4). ***p: <0.001 with 2-way ANOVA Tukey’s multiple comparisons test. **(b)** Overall experimental set up for the phosphoproteome analysis of human myotubes with perturbed CDK1 activity and expression**. (c)** Heatmap of phosphorylation changes (log2 fold changes) in the CDK1 target sites at 5 min and 30 min following insulin stimulation. The cells with asterisk indicate statistically significant changes at 5% FDR (red and blue indicating increase and decrease from 0 min, respectively). Gray cells indicate the absence of data points.

The pharmacological inhibition made a greater impact on the temporal phosphorylation profiles on the CDK1 targets than the siRNA-mediated knockdown overall (**Figure 4c**). First, the phosphorylation of multiple residues in ribosomal protein S6, a 40S ribosomal subunit, increased upon insulin stimulation only in CDK1 inhibited cells but not in cells with siRNA knockdown. Likewise, canonical phosphorylation on EIF4B, laminin A (LMNA), serine and arginine rich splicing factor 1 (SRSF1), pre-mRNA processing factor 4B (PRPF4B), and tuberin (TSC2) were attenuated in CDK1 inhibited cells compared to their respective controls. The inhibitory phosphorylation events on AKT1S1 (PRAS40) were also more active in the myotubes treated with the CDK1 inhibitor, rather than the siRNA knockdown. The phosphorylation levels of 5’-AMP-activated protein kinase subunit beta-1 (PRKAB1), general transcription factor IIi (GTF2I), heparin binding growth factor (HDGF), nucleolin (NCL), and interferon regulatory factor 2 binding protein 2 (IRF2BP2) were more dynamically regulated in the CDK1 inhibited cells, not in the cells with siRNA knockdown.

More importantly, the aforementioned phosphorylation events on EIF4B, SRSF1, LMNA, PRPF4B, and TSC2 increased upon insulin stimulation in the control cells with normal CDK1 activity (DMSO) only, but not in the CDK1 inhibited cells. Further, even though it was only measured in one of the three subjects, both CDK1 inhibition and siRNA knockdown desensitized the inhibitory phosphorylation signaling of AS160 (TBC1D4) for its GTP-ase activity that triggers GLUT4 membrane translocation^34^, whereas its regulatory phosphorylation was present in the respective control cells. Another interesting observation was the markedly reduced serine 1101 phosphorylation on IRS1 reproducibly observed in the CDK1 inhibited samples from two samples of Chinese donors. Combined with the fact that this site is a direct target site of P70S6K (RPS6KB1) ^35^, we hypothesize that the lack of CDK1 activity also desensitizes one of the potential feedback mechanisms of the mTORC1 signaling cascade to the IRS1 activity in myotubes.

## Discussion

In this work, we described the shared and unique proteome and phosphoproteome changes in response to a mild dose of insulin in human myotubes collected from three groups of ethnic background and distinct obesity status. These changes spanned across cellular processes in and out of insulin signaling in myotubes, and they showed differences across the three groups. We remark that our investigation was based on the myotubes derived from biopsy-isolated primary myoblasts following a protocol to minimize cellular heterogeneity from biopsies. To the best of our knowledge, this study has the largest number of human myotubes (N=30) with global and dynamic proteome and phosphoproteome data. By studying the insulin stimulated signaling cascade in subjects without type 2 diabetes, we also avoided confounding from abnormal metabolic activities in participants with hyperglycemia or undergoing treatment. Meanwhile, recruitment of subjects from different BMI and ethnicity backgrounds allowed us to study the inherent metabolic characteristics of different donors and compare their molecular profile in response to insulin stimulation.

Our data portray a modest yet acute shift of the myotube proteome and phosphoproteome in response to insulin stimulation (5 min), followed by persistent changes in key insulin signaling agents that occur over a longer time span in the phosphoproteome (30 min). Consistent with previous studies in muscle fibers ^7^ and stem cell-derived myoblasts ^6^, the changes we observed encompassed biological processes far beyond insulin signaling. We have recaptured protein abundance and/or phosphorylation changes expected to occur in the key proteins within the insulin signaling pathway such as IRS2, AKT, SOS, TSC1, RPS6, AS160, PIK3CA and mTOR. Outside insulin signaling, the dynamic changes included the proteins annotated to be part of mRNA splicing, signaling with Rho GTPase, tight junction and growth factor signaling pathways. Moreover, our KSA2D analysis enabled direct exploration of the correlated changes between kinase abundance levels at 5 min and substrate phosphorylation levels at 5 min and 30 min. In addition to the expected increase in kinases activity of AKT, MAPK1, MAK3, PRKACA, PRKCD, we discovered novel hub kinases such as serine/threonine protein kinase PAK1 and CDK1.

### Phosphorylation changes in canonical insulin signaling cascade

IRS1 and IRS2 are the first-line upstream responders to insulin signal cue transduced by the transmembrane receptor INSR. In our study, acute increase in IRS2 abundance at 5 min was observed across the thirty subjects. Independent of the protein abundance changes, the phosphorylation level of IRS2 (S608) increased at 30 min. Batista *et al*. discussed that increased serine phosphorylation of IRS2 has been linked to overactivity of a variety of serine/threonine kinases, including novel isoforms of PKCs and S6K, and this was replicated in our data through KSEA. Moreover, phosphorylation of AKT2 (T309), the equivalent event of AKT1 (T308) by PDK1, is known to be crucial in activating downstream substrates in insulin signaling. Previous studies have shown a decreased phosphorylation level at this residue in T2D patients ^36, 37^. Its activation is captured in our data as we observed increased phosphorylation at both 5 min and 30 min post stimulation, verified through western blots. Regulated activation of IRS2 and AKT2 reflects canonical insulin response in our myotube samples.

AS160 (TBC1D4) is a major AKT substrate. Once phosphorylated, its basal role as a Rab GTPase-activating protein (Rab GAP) that switches Rab proteins from the GTP-bound form to the inactive GDP-bound form is inhibited ^38^. Therefore, phosphorylated AS160 increases the abundance of the GTP-bound Rabs, which then form vesicles containing GLUT4 and fuse them into the cell membrane to facilitate glucose uptake. Phosphorylation of AS160 is required for GLUT translocation: point mutations of S318A, S588A, T642A and S751A disabled phosphorylation and resulted in 80% reduction of insulin-stimulated GLUT4 translocation ^39^. Our data showed increased phosphorylation levels of multiple residues (S330, S341, S666) on AS160 at 30 min. Our KSA2D analysis also verified the concordantly increased phosphorylation level of S588 at 5 min and 30 min, as well as those of S341 and S665 at 30 min along with increased AKT abundance at 5 min. The data suggests that our model recapitulates direct insulin signal as it pertains to glucose uptake.

### Signal transduction specific to obesity status and ethnic groups

Modest effect sizes we observed from the global proteome and phosphoproteome data reflect the heterogeneity of molecular events between groups and across different biopsy donors. Accordingly, the changes in the signaling events within some pathways were uniquely observed in certain obesity and ethnicity groups only. For example, serine phosphorylation on the death promoter gene BAD (S134) seems to be more pronounced in the obese group (CO) than in the lean group (CL), and in the South Asian lean group (SL) than in the Chinese lean group (CL) simultaneously. Inactive phosphorylation on this site in CO at both time points and in SL at 5 min likely reflects promotion of programmed cell death in these cells.

It appears that ethnicity as a factor makes a bigger difference in the global proteome and phosphoproteome than obesity status. One of the most noticeable differences between the two ethnic groups is that the key insulin signal receptor IRS2 has significant increase in the phosphorylation level in SL at 30 min compared to CL, specifically on S608. Second to this, we noticed milder phosphorylation changes on actin cytoskeleton elements at 30 min in SL, best exemplified by the decreased protein abundance of the members of actin cytoskeleton pathway including ACTN4, MYH9, CFL1 and TMSB4X. For example, ACTN4 has a role in tethering GLUT4 vesicles to the cortical actin cytoskeleton ^40^, and MYH9 is involved in insulin-stimulated docking and fusion of GLUT4 storage vesicle to the plasma membrane ^41^. A dysfunctional reorganization of the actin cytoskeleton has also been observed in insulin-resistant human and rodent skeletal muscles ^42^. However, how signalling pathways interconnect with the cytoskeletal elements remains relatively unexplored. Our study suggests that a dampened GLUT4 translocation and glucose uptake may contribute to insulin resistance in SL.

### KSA2D analysis reveals novel kinase hubs involved in insulin signaling

Our 2D analysis provides another unique angle for detecting kinase-substrate pairs that co-vary within the same individuals, including two kinases PAK1 and CDK1 that have rarely been implicated in the insulin-induced signaling pathway. PAK1 facilitates GLUT4 translocation to the plasma membrane, which is linked to its mediating role of the aforementioned Rac1 of Rho GTPase family ^43^. In skeletal muscle, Rac1 signals to PAK1 and facilitates its phosphorylation in response to insulin ^26^. PAK1 signals cofilin dephosphorylation and evokes F-actin remodeling, which is a prerequisite for GLUT4 vesicle translocation and glucose uptake into mature skeletal muscle. The signaling cascade is independent of the well-studied PI3K-AKT-AS160-Rab GTPase pathway, through which glucose uptake takes place ^26^. Previously, PAK1 has been shown to phosphorylate BAD that causes reduction of the physical interaction between BAD and BCL2, thus inhibiting pro-apoptotic effects of BAD ^44^. Similar to AKT1, the increased abundance of PAK1 at 5 min was well correlated with the increased phosphorylation level of various serine residues on BAD at 30 min (S99, S118, S134) (**Figure 3, Supplementary Table 6**). In addition to assisting glucose uptake, PAK1 might also promote cell survival and proliferation through its interaction with BAD. Nonetheless, BAD is only one of numerous substrates of PAK1 with increased phosphorylation levels. As such, the effect of its phosphorylation activity on the protein filaments of the cytoskeleton, such as stathmin (STMN1) and filamin A (FLNA) may also be worth investigating further.

CDK1 interacts with a broad spectrum of substrates, and its activity is especially pronounced in the SL group according to our data (**Supplementary Figure 6**). The canonical role of CDK1 in cell cycle regulation is well established, and it was therefore surprising CDK1 emerged as a key candidate kinase in our data. Unlike myoblasts, myotubes are terminally differentiated cells which has exited from the cell cycle. Within 72 hours of myogenic differentiation *in vitro*, myoblasts have been shown to undergoes a well-characterized sequenced of morphological and transcriptional changes to form post-mitotic plurinucleated myotubes ^45^. Our myotube characterizations, at day 7 post-differentiation, (**Supplementary Figure 2)** suggest that the myotubes lines used in this study are near or terminally differentiated. CDK1 was recently shown to be essential for myoblast proliferation, muscle regeneration and muscle fiber hypertrophy in satellite cell type specific Cdk1 knockout mice ^46^. However, its contribution to metabolic actions of insulin remains understudied, except a few examples such as enhancing glucose sensing in quiescent pancreatic beta-cell ^47^. Our findings suggest that insulin uses components of the cell cycle machinery in post-mitotic cells to control glucose metabolism independently of cell division. There is indeed increasing evidence for additional functions of cyclins and CDKs outside of their canonical roles ^48^. For instance, cyclin D1-Cdk4, has been shown to control glucose metabolism independently of cell cycle progression ^49^.

### Extra-mitotic role of CDK1 in human myotubes

An extensive analysis of the extra-mitotic actions of CDK1 was recently described by Haneke *et al*., illustrating that the kinase controls 5’TOP mRNA translation in a LARP1-dependent manner ^35^. In our study, CDK1 inhibition reduced the phosphorylation of LARP1 at S766 (**Figure 4c)**. Intriguingly, this residue is near a rapamycin-sensitive site at the C-terminal of LARP1 (T768). LARP1 is a key repressor of TOP mRNA translation ^50^ and mTORC1 promotes TOP mRNA translation through site specific phosphorylation of LARP1 ^51^. Additionally, mTORC1 also controls translation via direct regulation of S6K1 and 4E-BP1 ^52^. S6K1 (RPS6KB1) activates translation initiation through direct phosphorylation of eIF4B. The mTOR/PI3K and MAPK pathways involved in translational control converge to phosphorylate eIF4B on S422 ^53^. We demonstrate here that phosphorylation levels at S422 and S406 at eIF4B were reduced with CDK1 inhibition in all three subjects. In addition, RPS6 downstream of S6K1, was sensitive to CDK1 inhibition, especially on S244. Finally, phosphorylation level of S1101 residue of IRS1 was reduced with CDK1 inhibition. S1101 is a S6K1-specifc target ^54, 55^ and its phosphorylation negatively regulates PI3K/AKT signalling to induce insulin resistance ^56^.

mTORC1 also phosphorylates 4E-BP1 at multiple sites, causing the dissociation of 4E-BP1 from eIF4E to promote protein translation downstream. During mitotic entry, CDK1 can subsume the role of mTORC1 to phosphorylate some of mTORC1 substrates, including 4E-BP1 ^57^. Both mTORC1 and CDK1 phosphorylate 4E-BP1 at identical residues with CDK1 additionally phosphorylates 4E-BP1 at S82 in mice (S83 in humans). However, inhibition of CDK1 did not seem to affect 4E-BP1 in our study. Overall, the entry point of CDK1 in modulating the insulin signaling pathway is through multiple kinases, especially those involved in translation control via the mTORC1-S6K axis **(Figure 5)**, as expected in the landscape of myogenic and post-myogenic kinome ^21^.

**Figure 5.**
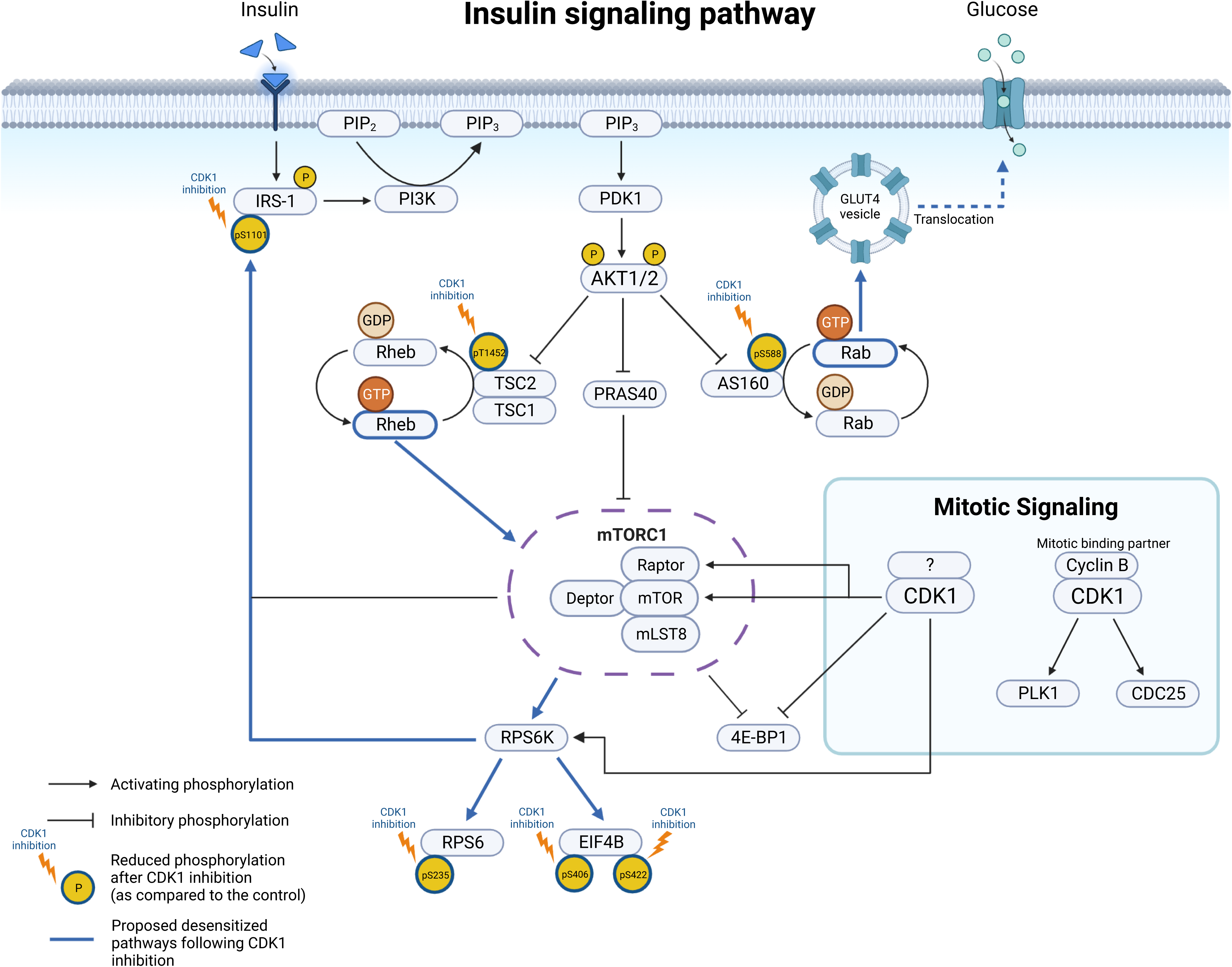
Effects of CDK1 inhibition on the insulin signaling pathway in primary human myotubes observed in the phosphoproteome analysis. The presumed basal extramitotic CDK1 activity mainly affects the mTORC1-S6K signaling arm of the canonical insulin signaling pathway. The exact mechanism of its involvement can be further investigated through the identification of potential effectors and substrate mapping in a greater number of myotube samples.

## Conclusion

Our study provides a rich resource to study the role of additional kinases in insulin signalling within human myotubes. However, there are several limitations that should be noted. First, while cultured human skeletal muscle cells maintained most metabolic properties in derived cell cultures, our observations are confined to myotubes and could not fully recapitulate myofibers^58^ *in vivo*. For instance, muscle fiber type composition varies between different regions of the body and the ratio of GLUT1:GLUT4 is reported to be higher in human myotubes compared with adult skeletal muscle ^59, 60^. Signalling defects associated with insulin resistance are also modified by gender and have functional differences in pathways downstream ^61^. However, all myotubes used in this study are derived from male donors and limits the validity of our findings on females. Finally, while our study revealed an extramitotic role of CDK1 in differentiated myotubes, this does not exclude the possibility that CDK1 has divergent roles in sub-populations of muscle cells ^62^. A small percentage of proliferating myoblasts do not differentiate and return to quiescence, or “alerted” states ^63^. Quiescent cells that transition into the alerted state possess enhanced tissue regenerative function as an adaptive response to injury and stress, priming them for cell cycle entry ^64^. However, we did not investigate the extent of cell heterogeneity within our study.

There are noticeable differences in signalling cascades between the two ethnic groups which require further investigations. IRS2 has significant increase in phosphorylation levels in SL compared to CL, specifically on S608, a residue which lies within the kinase regulatory-loop binding region (residues 591-733) that interacts with the tyrosine kinase domain of the insulin receptor and is dependent upon phosphorylation of the kinase activation loop ^65^. While IRS1 is required for myoblast differentiation and glucose metabolism, IRS2 is important for lipid metabolism and ERK activation in skeletal muscles ^66^. In addition, KSA2D analysis showed that phosphorylation on the substrates of CDK1 were more pronounced in SL. It is tempting to speculate that some of the underlying differences between South Asians and Chinese may be explained by disruption in the interplay between IRS1 and IRS2 in respond to nutrients cues and that CDK1 is implicated in regulating this. However, further experiments, especially in different nutrient status ^67, 68^ or with insulin resistance inducing perturbations, such as those recently investigated in adipocytes and adipose tissue ^69^, are required to examine this hypothesis and our understanding of CDK1 extramitotic role. Together with the differences noted in the cytoskeleton, our study reveals potential candidates which may contribute to ethnic differences in insulin-stimulated glucose uptake.

In summary, our data delineates the complexity of the connection between insulin signaling with other fundamental biological processes such as protein synthesis, cell fate decisions, actin cytoskeleton regulation, and cell growth. The cellular response of myotubes to insulin stimulation also has intrinsic connection to the ongoing debate whether the growth factor signaling can trigger myocyte proliferation by active cell division and how such connections can play a role in the context of the development of IR in healthy individuals. In addition, we also demonstrate that CDK1 may play a central role in connecting proliferative cues with protein synthesis through mTOR1C-S6K1 axis, and there may be additional kinases and phosphatases that may precondition efficient insulin signaling in these cells.

## STAR * Methods

### Subjects and clinical measurements

The current study is a sub-study of the Singapore Adult Metabolism Study (SAMs). SAMs is a cross-sectional study that aims to investigate the influence of ethnicity on metabolic disease in healthy, overweight and obese subjects. A total of 101 Chinese, 82 Malay and 81 South Asian males were recruited for SAMs. The study has been registered at clinicaltrials.gov as NCT00988819. A sub-set comprising 30 subjects, 10 each from CO (body mass index (BMI) >25.0 kg/m^2^), CL and SL (BMI 19-24.0 kg/m^2^) were chosen for this specific study. All subjects had fasting blood glucose of < 7mmol/L. Exclusion criteria briefly includes current smoking and/or prior history of hypertension, dyslipidemia, heart disease, epilepsy, insulin allergy, ingestion of drugs reported to affect insulin sensitivity, hospitalization and surgery 6 months before enrolment into the study. Further information on the SAMs recruitment process and subject details are as previously described (4). Ethics approval was obtained from the National Healthcare Group Domain Specific Review Board (Singapore) (DSRB ref no. C/2009/00022). All subjects provided informed consent. Ethnicity, age, medical and drug history, and data on lifestyle factors were self-reported by the subjects and collected using interviewer administered questionnaires. Measurements of other clinical characteristics, including the height, weight, blood biochemistry panels and insulin sensitivity of the subjects are carried out as previously described (4). The collated clinical data characteristics are shown in **Table 1**.

**Table 1.** Clinical data summary

| Clinical characteristics | Chinese Obese | Chinese Lean | South Asian Lean | *P* value |
| --- | --- | --- | --- | --- |
| *N* | 10 | 10 | 10 | - |
| Age (years) | 32.7 (6.2) <sup>†</sup> | 28.4 (6.6) <sup>‡</sup> | 22.5 (1.8) <sup>†, ‡</sup> | <0.001* |
| Height (cm) | 174.1 (6.6) | 172.5 (5.1) | 176.4 (7.6) | 0.426 |
| Weight (kg) | 82.4 (8.4) <sup>†, ‡</sup> | 61.0 (5.3) <sup>†</sup> | 65.7 (7.2) <sup>‡</sup> | <0.001 |
| BMI (kg/m<sup>2</sup>) | 27.2 (1.5) <sup>†, ‡</sup> | 20.7 (1.1) <sup>†</sup> | 21.0 (1.1) <sup>‡</sup> | <0.001 |
| Waist circumference (cm) | 93.1 (4.6) <sup>†, ‡</sup> | 75.7 (4.9) <sup>†</sup> | 75.8 (4.6) <sup>‡</sup> | <0.001 |
| Hip circumference (cm) | 103.2 (3.4) <sup>†, ‡</sup> | 93.3 (5.0) <sup>†</sup> | 93.8 (4.7) <sup>‡</sup> | <0.001 |
| Fasting glucose (mmol/L) | 4.68 (0.44) | 4.47 (0.43) | 4.39 (0.25) | 0.216 |
| Fasting insulin (pmol/L) | 11.42 (5.61) <sup>†</sup> | 5.88 (1.84) <sup>†</sup> | 7.72 (3.32) | 0.012 |
| Total cholesterol (mmol/L) | 5.48 (1.21) <sup>†</sup> | 4.50 (0.71) | 4.02 (0.67) <sup>†</sup> | 0.004 |
| LDL cholesterol (mmol/L) | 3.62 (1.12) <sup>†</sup> | 2.77 (0.60) | 2.42 (0.51) <sup>†</sup> | 0.007 |
| HDL cholesterol (mmol/L) | 1.22 (0.22) | 1.39 (0.28) | 1.27 (0.22) | 0.313 |
| Triglycerides (mmol/L) | 1.39 (0.68) <sup>†, ‡</sup> | 0.76 (0.27) <sup>†</sup> | 0.72 (0.14) <sup>‡</sup> | 0.028* |
| % Body fat (%) | 26.3 (4.4) <sup>†, ‡</sup> | 16.8 (3.6) <sup>†</sup> | 18.0 (5.1) <sup>‡</sup> | <0.001 |
| ISI (mg/kg lean mass) | 8.2 (4.3) <sup>†</sup> | 15.5 (6.3) <sup>†, ‡</sup> | 9.7 (1.7) <sup>‡</sup> | 0.029* |

Data are presented as means (SD) with *P* values from ANOVA tests across the three groups. Abbreviation: ISI insulin sensitivity index
\**P* value derived from Welch's ANOVA test due to unequal variances.
<sup>†, ‡</sup> Significant Tukey's and Games-Howell's post hoc test results for pairwise comparison among the groups with equal and unequal variances respectively (*P* value <0.05).

### Cell culture and differentiation

Primary human myoblast samples were isolated from skeletal muscle biopsies of the 30 subjects using protocol previously described by Khoo et al ^70^. The myoblasts were cultured on 10% Matrigel-coated dishes (Corning, 354234) and maintained in growth media containing Dulbecco’s Modified Eagle’s Medium (DMEM) (Gibco, Thermo Fisher Scientific, 11995) supplemented with 20% Fetal Bovine Serum (Hyclone, GE Healthcare Life Sciences, SV30160.03), 10% Horse Serum (HS) (Gibco, Thermo Fisher Scientific, 16050), 1% Penicillin and Streptomycin (P/S) (Hyclone, GE Healthcare Life Sciences, SV30010) and 1% Chicken Embryo Extract (MP Biomedicals, 2850145). Medium was changed every other day. Each myoblast sample was seeded on 150 mm culture dishes at an initial density of 0.8 × 10^6^ and were slowly expanded to 15 150 mm dishes. The myoblasts were grown to confluence before initiation of differentiation. Myogenic differentiation was achieved with incubation in differentiation media consisting of DMEM, 2% of HS and 1% P/S for 7 days ^71^. All cells were grown at 37 °C and 5% CO_2_ in a humidified environment.

### Insulin stimulation and harvesting of myotubes

On day 7 of differentiation, the culture media was replaced with serum-free minimum essential medium-alpha (Gibco, Thermo Fisher Scientific, 12561). The medium was removed 14 hours later and the cells were washed with phosphate buffered saline (PBS). The myotubes were then treated with (1,5,15 and 30 minutes) or without (control, 0 minutes) 10 nM of insulin (Gibco, Thermo Fisher Scientific, 12585) at 37°C to sufficiently stimulate, but not saturate insulin signaling in the myotubes. At each stipulated time-point, insulin-containing media was removed, washed with PBS and TrypLE Express Enzyme (Gibco, Thermo Fisher Scientific, 12605) was added for 5 to 10 minutes to detach the myotubes from the dishes. The dishes were also scraped with a cell lifter to ensure maximum harvest of the cells. The collected myotubes were centrifuged at 2200*g* for 10 minutes (Sorvall Legend X1R, Thermo Fisher Scientific) and the supernatant was discarded. The resulting cell pellets were frozen and stored at -80°C for use in downstream studies.

### Phosphorylated AKT Immunoblot Analysis

The insulin response of the treated cells was evaluated using the insulin-dependent AKT phosphorylation (serine residue 473) as an index of myotube insulin sensitivity. The cell pellets were first lysed in RIPA buffer supplemented with EDTA and a protease and phosphatase inhibitor cocktail (Thermo Fisher Scientific). Protein concentration of the cell lysates were determined with Bradford assay (Bio-Rad). Equal amounts of protein were mixed with Laemmli sample buffer (Bio-rad) and B-mercaptoethanol (Bio-rad) as per manufacturer’s instruction, before being subjected to 100 °C heat treatment for 5 minutes. Pre-stained protein ladder (Abcam, ab116028) was used to estimate the approximate molecular size of bands. Samples were run on 4-20% Mini-PROTEAN TGX precast gels (Bio-Rad) in Tris/Glycine/SDS running buffer (Bio-rad). The proteins were then transferred onto a 0.2 μm PVDF membrane. The membranes were blocked with StartingBlock T20 (TBS) Blocking Buffer (Thermo Fisher Scientific) and incubated overnight at 4 °C with primary antibodies raised against phospho-AKT Ser473 (Cell Signaling Technology, 9271S), total AKT (Cell Signaling Technology, 9272S) and GAPDH (Cell Signaling Technology, 5174S). The membranes were then washed with PBS-Tween and incubated with anti-rabbit antibodies conjugated with horseradish peroxidase (Cell Signaling Technology, 7074). After a final round of washing, the membranes were incubated with Clarity ECL substrate (Bio-Rad) and exposed on Tru-Pro XBT ECL Exposure Films (Research Instruments). Band intensities were quantified using ImageJ analysis and normalised to GAPDH.

### RNA isolation, cDNA synthesis and Quantitative Polymerase Chain Reaction (qPCR)

Total RNA was extracted from cultured human primary myoblasts (Day 0) and myotubes (Day 7) using the ISOLATE II RNA Mini kit (Bioline). Isolated RNA were then reverse transcribed using iScript™ Reverse Transcription Supermix (Bio-rad). The resulting cDNA was used to perform qPCR analysis with the iTaq Universal SYBR Green Supermix (Bio-rad) on an ABI Vii7 instrument (Applied Biosystems). All processes were carried out as per each manufacturer’s instructions. The primers for each target gene are listed in **Table 2**. The data was analysed using the comparative Ct method to calculate relative gene expression normalised to RPLP0.

**Table 2.**
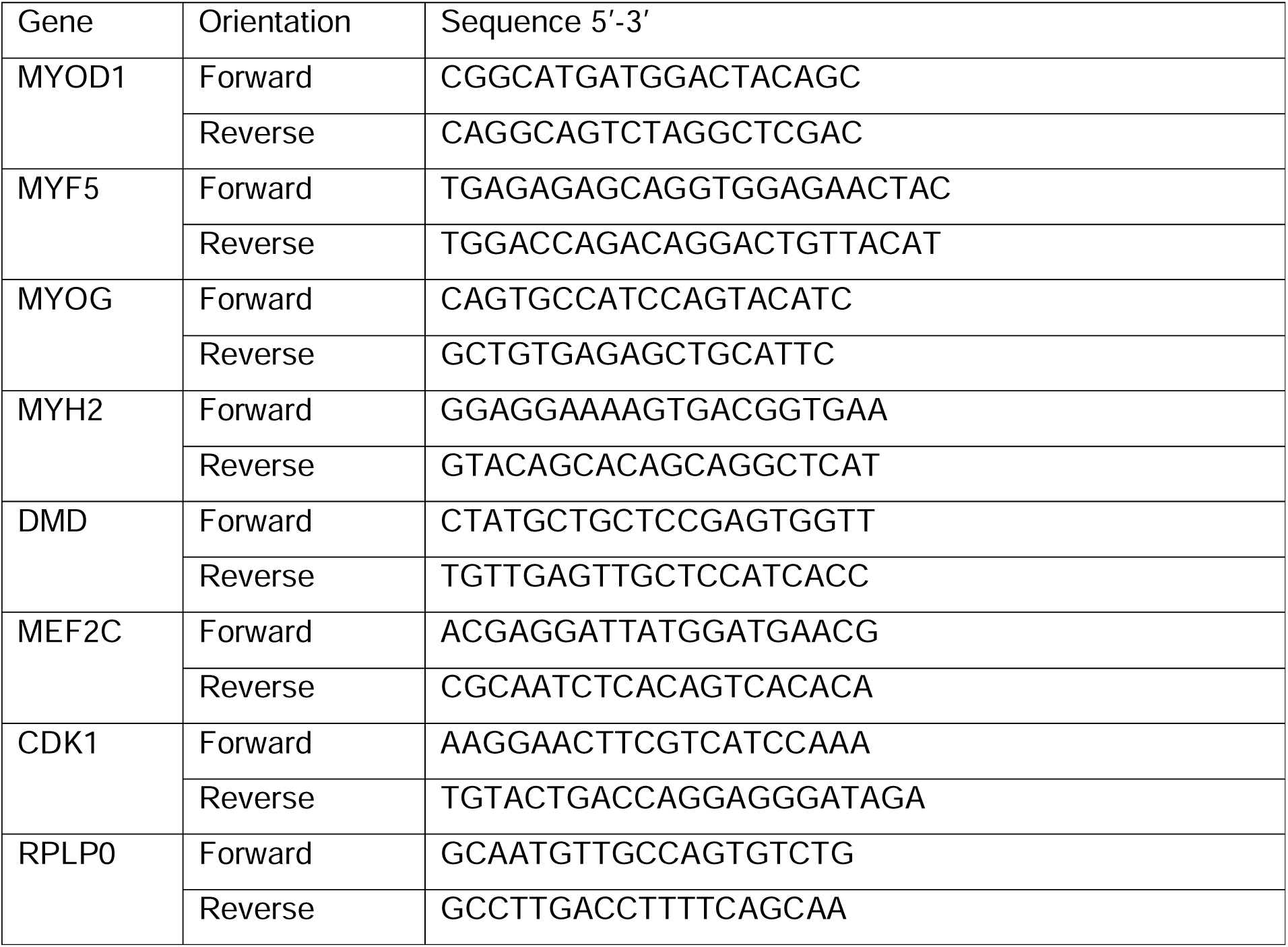
List of qPCR primers used for gene expression analysis

### Immunofluorescence staining

Human primary myoblasts were seeded and grown on 12-well plates. Growth and differentiation media were then removed from the myoblasts (Day 0) and myotubes (Day 7) respectively. The cells were fixed with 10% formalin (Sigma Aldrich), permeabilised with 0.1% Triton-X (Sigma Aldrich) and blocked with 10% FBS in PBST for 30 minutes. Plates were subjected to washes with PBS in between steps. Cells were stained for 1 hour at room temperature with the following primary antibodies and concentrations: Desmin (Abcam, ab15200, 1:300) and Fast Myosin Skeletal Heavy Chain (Abcam, ab51263, 1:200). The following secondary antibodies were used with the non-conjugated primary antibodies: Goat Anti-Mouse IgG H&L Alexa Fluor 488 (Abcam, ab150113, 1:200) and Rabbit IgG H&L Alexa Fluor 555 (Abcam, ab150078, 1:500). Lastly, 4′, 6-diamidino-2-phenylindole (DAPI) (Invitrogen, D1306) was used as a nuclear counterstain at a final concentration of 1.43μM. Images were captured with an Olympus FluoView FV3000 (Olympus, Japan) laser scanning confocal microscope using a 10x/0.40 objective. Digital zoom at 2X and 5X were done using the Olympus Fluoview software.

### Glucose uptake assay

Glucose uptake assay was done using the Glucose Uptake-Glo^TM^ Assay kit (Promega, J1343). On day 7 of differentiation, culture media was changed to serum free DMEM (Gibco, Thermo Fisher Scientific, 11995). The medium was removed 14 hours later, and the cells were washed with PBS. Subsequent steps were done according to manufacturer’s protocol, with brief modifications. Briefly, the cells were treated with 1 µM of insulin in glucose free media (Gibco, Thermo Fisher Scientific, 11966) for 1 hour at 37 °C in 5% CO_2_. After incubation, the insulin containing media was removed and replaced with 1mM of 2-deoxyglucose in PBS and incubated for 10 minutes at 25 °C. Stop buffer was added after to lyse the cells and stop all reactions. Neutralisation and detection reagent were then added and the resulting solution was incubated for 1 hour at 25 °C. Luminescence reading was done on Varioskan Flash microplate reader (Thermo Fisher Scientific).

### CDK1 inhibition and CDK1 siRNA mediated knockdown

#### CDK1 inhibition using RO-3306

CDK1 inhibition was carried out using a pharmacological inhibitor, RO-3306 (Sigma Aldrich, SML0569). RO-3306 was reconstituted in dimethyl sulfoxide (DMSO) to a stock concentration of 10mM and kept frozen at -20 °C. Myoblasts were cultured and differentiated as per the protocol stated above. On day 8 of differentiation, the culture media was replaced with either 0 μM (DMSO) or 5 μM of RO-3306 in serum-free DMEM (Gibco, Thermo Fisher Scientific, 11995) and incubated for 3 hours. Following the 3 hours serum free DMEM incubation, subsequent steps for glucose uptake, insulin stimulation and harvesting of myotubes are as described above with the addition of DMSO or 5 μM RO-3306 throughout the steps to ensure that constant inhibition of CDK1 activity.

#### CDK1 siRNA mediated knockdown

All siRNA knockdown reagents were purchased from Dharmacon. ON-TARGETplus Human CDK1 SMARTpool (L-003224-00-0005) and ON-TARGETplus non-targeting pool siRNAs (D-001810-10-05) were reconstituted in siRNA buffer (B-002000-UB-100) to a concentration of 20 μM and kept frozen in -20 °C. Myoblasts were cultured and differentiated as per the protocol stated above. On day 4 of differentiation, the cells were transfected with either 25 nM of non-targeting siRNAs or CDK1 siRNAs using DharmaFECT 1 transfection reagent (T-2001-03). Transfection was performed according to the protocol provided by the manufacturer. After 96 hours, the transfection mix was removed and replaced with serum free DMEM (Gibco, Thermo Fisher Scientific, 11995) and further incubated for 3 hours. All subsequent steps for glucose uptake, insulin stimulation and harvesting of myotubes are as described above.

### Proteome and phosphoproteome analysis

Protein extraction and tryptic digestion. Myoblasts derived from 10 Chinese obese, 10 Chinese lean and 10 South Asian lean subjects were lysed in 600 µL of lysis buffer (Thermo RIPA buffer, 89900, 534 µL, 100x Halt™ Protease Inhibitor Cocktail, 78430, 6 µL, 10x Roche PhosSTOP™, 4906845001, 60uL) using probe homogenizer. Cell lysate was centrifuged at 16,000g, 4 °C for 10 minutes, and the protein concentration of each sample was determined by BCA assay (Micro BCA™ Protein Assay Kit, Thermo Fisher Scientific, 23235). Each 500 µg of protein was digested individually using a filter assisted sample preparation (FASP) method ^72^. A universal standard was used to correct measurement errors across experimental batches, which was prepared by pooling in the same amount from all 90 samples (Figure 1b). Proteins of each sample were denatured and reduced in SDT buffer (4% sodium dodecyl sulfate (SDS) in 0.1 M Tris-HCl, pH 7.6 and 0.1 M Dithiothreitol (DTT)) at 37 °C for 1 hour with shaking and boiling for 10 minutes at 100 °C. The buffer was changed to 6 M urea in 0.1 M Tris-HCl, pH 8.5 and the proteins were alkylated with 0.5 M iodoacetamide (IAA) for 30 minutes at room temperature in the dark. 96 samples (90 individual subjects and universal standards for 6 experimental batches) were subjected to digestion with LysC enzyme (Wako Chemicals, 121-05063) at 1 mAU:50 enzyme to substrate ratio for 2 hours at room temperature (RT) followed by the MS grade trypsin (Pierce Biotechnology, 90057) at a 1:50 enzyme to substrate ratio and overnight incubation at 37 °C. The resulting tryptic peptides were dried using Speed-Vac (Hypercool; Labex) and kept in -80 °C until the tandem mass isobaric tag (TMT, Thermo Fisher Scientific, A44520) labeling.

#### TMT-labeling of peptides and basic pH reverse phase fractionation

Peptides were labeled with 16 different mass tags among the Tandem Mass Tag (TMT) pro™ 16plex reagents (Thermo Fisher Scientific, A44520) as follows: Samples corresponding to each of the 3 time points of 5 patients were included per experimental batch (126N to 133C), and a total of 6 sets of labeling were performed. The last TMT channel (134N) for each batch is a universal standard and was used for normalization of experimental batches (**Supplemental Table 1)**. The prepared TMT reagent was transferred to the peptide sample and the mixture was incubated for 1hr at room temperature following brief vortexing. The reaction was quenched with 5% hydroxylamine and incubate for 30 minutes at room temperature. All TMT labeled peptides in the batch were pooled and concentrated by vacuum centrifugation. Labeled peptides were loaded on an analytical column (XBridge Peptide BEH C18 Column, 300 Å, 5 µm, 4.6 mm × 250 mm, Waters™, 186003625) for fractionation. A gradient was generated using an Ultimate 3000 HPLC system (Dionex, Germany) operated with solvent A (4.5 mM ammonium formate (pH 10) in 2% (vol/vol) acetonitrile) and solvent B (4.5 mM ammonium formate (pH 10) in 90% (vol/vol) acetonitrile). The gradient was as follows: 0–7 min, 0% B; 7–13 min, 16% B; 13-73 min, 40% B; 73-77min, 44% B; 77-82min, 60% B; 82–96min, 60% B; 96–110min, 0% B ^73^. The separated peptides were collected and non-consecutively pooled into 12 fractions by combining 8 parts (#1, #13, #25, #37, #49, #61, #73, and #85; #2, #14, #26, #38, #50, #62, #74, and #86 and so on). The resulting 12 fractions were desalted with a C18 spin column. A 95% amount of the sample was subjected to phosphopeptide enrichment and the remaining amount of sample was used for global analysis (**Figure 1b**).

#### Phosphopeptide enrichment using Fe-IMAC

Before phosphopeptide enrichment, the 12 fractions samples were concatenated into 6 fractions by combining 2 parts (#1, #7; #2, #8; and so on). For enrichment, using immobilized metal affinity chromatography (IMAC) as previously described ^74^. In brief, Ni-NTA agarose bead slurry (QIAGEN, 36113) was washed with deionized water and then reacted with 100 mM EDTA (pH 8.0) by mixing for 30 minutes on an end-over-end rotator (SB3, Stuart, UK). The beads were then reacted with freshly prepared 10 mM aqueous FeCl_3_ solution for 30 minutes with end-over-end rotation. For each of the 6 fractions, peptides were reconstituted to 0.5 mg/mL in IMAC binding/wash buffer (80% acetonitrile, 0.1% formic acid) and incubated with the Fe^3+^-IMAC beads for 30 minutes at RT. After incubation, the beads were washed 2 times using IMAC wash buffer and eluted 500 mM dibasic potassium phosphate, pH 7.0 a total of three times. The eluted phosphopeptides were acidified immediately with 10% TFA to pH 3.5-4.0 before vacuum dry and for LC-MS/MS analysis, the enriched phosphopeptides were desalted using a C18 spin column (Thermo Fisher Scientific, 89870).

#### LC-MS/MS analysis for global proteome and phosphoproteome

For LC-MS analysis, Ultimate 3000 RSLC nano system (Dionex, Germany) was coupled to Q Exactive HF-X hybrid quadrupole-Orbitrap mass spectrometer (Thermo Fisher Scientific, Germany) and equipped with a trap column (Acclaim™ PepMap™ 100 C18 LC Column, C18, 75 µm × 2 cm, 5 μm particle size, Thermo Scientific Inc., 164564) for clean-up followed by an analytical column (EASY-Spray™ LC Columns, C18, 75 µm × 50 cm, 2 μm particle size, Thermo Scientific Inc., ES803A). The peptides were separated by using the mobile phase comprising of solvent A (0.1% formic acid and 2% acetonitrile in water) and solvent B (0.1% formic acid in 90% acetonitrile). For global proteome digested peptides, the optimized linear gradient elution program was set as follows: (Tmin / % of solvent B): 0/2, 6/2, 8/3, 103/30, 113/40, 114/90, 119/90, 120/2, 135/2 and the flow rate was 300 nl/min throughout the run time. For enriched phosphopeptides, the optimized linear gradient elution program was set as follows: (Tmin / % of solvent B): 0/2, 6/2, 8/3, 103/35, 113/40, 114/90, 119/90, 120/2, 135/2. Full MS scans were acquired for the mass range of m/z 350 – 1800 at the resolution of 120,000 in MS1 level, an automatic gain control (AGC) target of 3e6 and the MS/MS analysis was performed by data-dependant acquisition in positive mode. The analytical columns were maintained at 60 °C. The electric potential of electrospray ionization was set to 2.1 kV, and the temperature of the capillary was also set to 250 °C. The top 12 precursor peaks were selected from the MS1 scan and separated for fragmentation. In the MS/MS acquisition, the resolution was set to 45,000 with a fixed first m/z of 100 m/z, an AGC target of 2e5, an isolation window of 0.8 m/z, and a normalized collision energy of 32. Ions with unassigned charge state, 1, or >6 were discarded. In the case of the phosphoproteome, analysis was performed with the same MS parameters as the global proteome for up to 10 MS/MS spectra.

#### Protein database searching and quantification of global and phosphoproteomic data

The MS/MS spectra were searched against a composite database of UniProt homo sapiens reference (April, 2020; 20595 entries) using Sequest HT search engine. Trypsin specificity was determined at up to two missed cleavages. In modification, carbamidomethylation, alkylation of disulfide bonds in cysteine (+57.021 Da), TMT 16-plex modification of lysine, and N-termination (+304.313 Da) were noted as static modifications. In common with global and phosphoproteomics data, oxidation of methionine (+15.995 Da) was noted as a variable modification. Dynamic modification of phosphorylation of serine, threonine, and tyrosine (+79.980 Da) added parameters to phosphoproteomics data. The FDR for peptide level was evaluated as 0.01 for removing false positive data as much as possible. To quantify each reporter ion, “peptide and protein quantifier” method in Proteome Discoverer 2.4 (Thermo Fisher Scientific, USA) with TMT 16-plex was used. For highly confident quantifications of protein, the protein ratios were calculated from two or more unique quantitative peptides in each replicate.

### Statistical analysis and graphics

#### Data quality control and normalization

In the TMT proteomics and phosphoproteomics data, each of the thirty subjects has three time points, at 0 min, 5 min, and 30 min post the insulin stimulation. Using the 16-plex TMT labeling, this implies that all time points for five individuals were included in one experiment. Accordingly, all 90 samples (30 subjects at three time points) were distributed to six TMT sets. We examined the batch effect and confirmed that reporter ion intensities between samples in different batches were not comparable, nor can be corrected by computational means (**Supplementary Figure 1**). However, the time-course changes, represented by the fold changes at 5 min and 30 min relative to the 0 min (time of stimulation) were not subject to significant batch effects, and as such, we transformed both proteome and phosphoproteome data into those ratios computed within each subject. For phosphoproteome data, we first mapped the phosphopeptides to Uniprot protein sequences to get the positions of the modified residues and summarized the quantitative data by averaging phosphopeptide intensities at unique residue (S/T/Y) in each sample. For both proteome data and phosphoproteome data, we added a fudge factor equivalent to the 1 percentile point of total abundance/phosphorylation to attenuate the issue of over-estimated fold changes in proteins and peptides with near zero intensities.

#### Empirical Bayes analysis of differential expression (KSA-1D): 1D analysis

For the differential expression analysis in proteomics and phosphoproteomics data, we implemented an empirical Bayes method that had been well established in high dimensional data analysis ^75^. The first step in modeling the expression ratio between two time points is to compute Z statistics across all the samples for each protein or phosphorylation site. To be specific, for protein *i* ∈ {1,2, …, *M*} in sample *j* ∈ {1,2, . . ., *N*}, with the measurement at 0 min and 5 min in log scale being *A*_*ij*0_ and *A*_*ij*5_, then, the time point ratio between 5 min and 0 min in natural logarithmic scale can be written as *D*_*ij*5_ = *A_ij_*_5_ - *A_ij_*_0_, and we use *D_i_*_5_ to represent the vector of the ratios across all the samples.

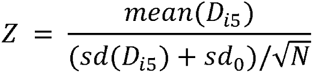

Here *sd_0_* is the 10th percentile point of all the standard deviations across all the proteins that is known to attenuate unorthodox behavior of the statistics in low abundance molecules ^76^. Throughout the analysis, we only used the proteins and phosphopeptides with missing data in fewer than 6 samples for construction of a reliable Z score. We also remark that, for phosphoproteome data, we normalize the phosphopeptide intensities by the protein-level intensity of the host protein in order to capture the net variation of phosphorylation events without the impact of whole protein abundance changes ^77^.

We model the distribution of all Z scores derived from all the proteins *f* as a mixture of proteins which are up- or down-regulated between time points and proteins which are non-differentially regulated. Assuming the Z scores of the differentially regulated proteins follow an empirical distribution *f*_l_ and the Z scores of the non-differentially regulated proteins follow an empirical null distribution, we can write the observed distribution for Z score as

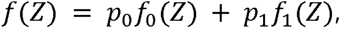

where *p*_0,_ *p*_1_are the proportion of the non-differentially and differentially regulated proteins in the data. We have *p*_0_ + *p*_l_ = 1 . To estimate the distribution of the non-differentially regulated proteins f_O_, we simulate the null distribution of Z scores by randomly permuting the measurements of the two time points for many times. Following this mixture deconvolution, we calculate the posterior probability of being differentially regulated for each protein with the following equation:

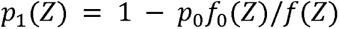

We estimate the prior probability p_O_ of being non-differentially regulated by comparing the density of estimated overall density and null density at *Z*_0_ = 0, *f(Z*_0_)/*f*_0_(*Z*_0_). Following Efron et al. ^75^, the local false discovery rate (lfdr) is computed as

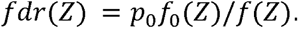

This analysis is called 1D analysis in the paper, which is applied to proteomics data and phosphoproteomics data separately. We present an extended 2D analysis method below for the analysis of known kinase-substrate pairs.

#### 2D analysis with kinase abundance and substrate phosphorylation (KSA2D)

Kinase and residue level substrate relationships were collected from three independent sources, including PhosphoSitePlus ^78^, PhosphoNetwork ^79^, and TyrosineKinase ^80^ for comprehensive coverage. After mapping the proteome data to kinases and the phosphoproteome data to substrates (with site level resolution), we gathered 2,122 kinase-substrate pairs for “2D” differential analysis, including 1,438 pairs recorded in PhosphoSitePlus, 765 in PhosphoNetwork, and 9 in TyrosineKinase.

Following the principles described in the 1D analysis, we derive two-dimensional Z score vectors for both kinase and substrate target site

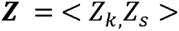

for each pair of kinase-substrate relationships and simulate the two-dimensional null distribution of Z scores by random permutations. *f* and *f*_0_ are estimated by the two-dimensional empirical density distribution, and the proportion of null *p*_0_is estimated by taking the ratio of densities at *Z*_00_= < *Z*_k_ =0, *Z_s_* =0 >, *p*_0_ = *f*(*Z*_00_)/*f*_0_(*Z*_00_). Similarly, we have,

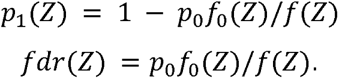

This analysis is called 2D analysis hereafter. The entire workflow, including data normalization, mapping from phosphopeptides to unique phosphorylation sites, and 1D/2D differential analysis are available in an R package KSA2D, freely downloadable from https://www.github.com/ginnyintifa/KSA2D.

#### Gene enrichment analysis

Upon the identification of significant up-regulated or down-regulated proteins and phosphorylation sites (local false discovery rate <0.01), we compare the list of genes encoding the significant proteins and phosphopeptides against the background list of all identified proteins using pathway enrichment analysis. Pathway information used was collected from ConsensusPathDB database ^81^. This is done using hypergeometric probability, i.e. the probability of having *k* out of *n* significant genes being in a certain pathway containing *K* genes is

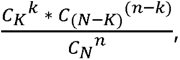

where N is the number of genes in the experiment, and the statistical significance score (*p*-value) is derived by adding up the probabilities of more extreme events. Significantly enriched pathways were determined at p<0.05.

#### Kinase-Substrate Enrichment Analysis (KSEA)

Kinase-Substrate Enrichment Analysis (KSEA) was conducted by the R package KSEAapp ^82^. The package takes the fold change of each phosphorylation site in our phosphoproteome data as the input and outputs scores for kinases based on the collective phosphorylation changes of their identified substrates.

#### Figure generation

Figures pertaining to statistical analysis were drafted using GraphPad Prism, R (version 4.3.1), Cytoscape (version 3.10.0) and Biorenderer.com and assembled into multi-panel figures using Adobe Illustrator (2023).

## SUPPLEMENTAL INFORMATION

Supplemental Information includes six figures and seven tables. They can be found online at http://URL_at_the_journal.

## AUTHOR CONTRIBUTIONS

CMK, MKL and YSL collected the clinical data and the biosamples from all study participants and designed the overall study with EST. KPK, MHL, and EST designed the experiments involving myotubes. BH and KPK performed untargeted proteomics and phosphoproteomic experiments for pilot and main analysis. EC, WLL, SH, and MHL performed myoblast isolation, myotube differentiation experiments, and western blot analysis. WLL and MHL performed CDK1 inhibition and siRNA knockdown experiments, and TZ, LCW and RS performed phosphoproteomics experiments for those samples. GXL and HC performed biostatistical and bioinformatics analysis. GXL, WLL, and BH drafted the manuscript, and GXL, BH, WLL, MHL, HC, KPK, and EST wrote the paper with input from all authors. All authors reviewed and provided input to the final manuscript.

## ACKNOWLEDGMENTS

The authors thank Prof. Philipp Kaldis for the guidance and helpful suggestions he provided during the data interpretation. This work was supported in part by grants of Korea Health Technology R&D Project through Korea Health Industry Development Institute (KHIDI), funded by the Ministry of Health & Welfare, Republic of Korea (HI15C1595 to K.P.K), and National Medical Research Council of Singapore, Cooperative Basic Research Grant (NMRC/BNIG14May019 to M.H.L), National Medical Research Council of Singapore (NMRC/CG/M009 to E.S.T), and Clinician Scientist Award from National Medical Research Council of Singapore (NMRC/CSA-SI/0002/2015 to E.S.T). The investigators also acknowledge Sir Peter Gluckman, Professor at the University of Auckland, for his advice and guidance in establishing the research program.

## DATA AVAILABILITY STATEMENT

The raw mass spectrometry data for the main proteome and phosphoproteome profiling experiments can be found in PRIDE database (https://ebi.ac.uk/pride/) with accession number PXD042870 (username, password 1Gum6wNH). The data for the CDK1 inhibition and siRNA knockdown experiments can be found in the jPOST database with accession number JPST002220 (URL: https://repository.jpostdb.org/preview/289540332649e7b6f03a1b, access key: 4335).

