## Supplemental Information for "Unbiased phosphoproteomics analysis unveils modulation of insulin signaling by extramitotic CDK1 kinase activity in human myotubes"

**Overlap analysis of dynamic myotube and phosphorylation sites and changes with published data sets**

Upon confirming successful terminal differentiation of myoblasts into myotubes, the cells were lysed, digested, and labelled with 16-plex tandem mass tags (TMT). The TMT-labelled samples were then combined and fractionated into twelve fractions for LC-MS/MS analysis. From the fractionated samples, 5% of the fractionated samples were used for the global proteome analysis and the other 95% of the fractionated samples were used for the phosphoproteome analysis following iron-chelate, ferric nitrilotriacetate (Fe-NTA) phosphopeptide enrichment (**Supplementary Figure 1**).

We first compared the phosphosites covered by our data and the dynamic trends of phosphorylation following insulin stimulation with published data sets. Batista *et al*. profiled phosphoproteome responses to a 100 nM insulin treatment for 10 minutes in stem cell-derived myoblasts (iMyos) in T2D patients and healthy controls. In another study, Hoffman *et al*. studied the impact of acute exercise on phosphoproteome in human muscle fibers. In order to conduct fair comparisons among the three studies, we limited the comparison of phosphoproteome to canonical proteins only (excluding isoforms), which resulted in 18,299 sites in our data, 20,440 sites in Batista *et al*., and 9,183 sites in Hoffman *et al*. Considering the variation in site localization of phosphosites from MS/MS data, we allowed up to +/-5 amino acid position shifts in matching the detected sites (of the same residue) between studies.

Comparing our myotube data with Batista *et al*., we found 49.6% of sites covered in our experiment had matching phosphorylation events in their data (51.9% in insulin signaling, 49.5% outside). Conversely, 51.2% of their sites had matching events in our data (56% in insulin signaling, 51.1% outside). In the comparison between our data and muscle fiber data in Hoffman *et al*., the overlap was much poorer: only 21.1% of sites we detected were covered by Hoffman *et al*. (29.6% in insulin signaling, 20.9% outside), and 38.4% of the sites detected in their work were covered by our data (43.8% in insulin signaling, 29.5% outside). Batista *et al*. and Hoffman *et al*. also had a poor overlap at the site level: only 17.4% of the detected sites in Batista *et al*. appeared in Hoffman *et al*. (26.9% in insulin signaling, 17.1% outside), whereas 30% of the detected sites in Hoffman *et al*. appeared in the data by Batista *et al*. (41.9% in insulin signaling, 29.6% outside). In summary, the phosphoproteome data reported by the three studies have a moderate overlap in general, with a better overlap between our myotube data and myoblast data in Batista *et al*. and with a better overlap within the insulin signaling pathway than outside. Although the differences in cell composition and purity of starting materials are a primary source of moderate overlaps, another share of the discrepancies are perhaps attributable to the differences in phosphopeptide enrichment methods and the innate run-to-run variability of the untargeted LC-MS/MS analysis.

We next examined the correlation of fold changes between insulin treated versus untreated samples between our myotube data and the iMyos data (statistically significant sites in the latter, N=646). The Pearson correlation of the log2 fold changes at 30 min from 0 min were 0.30 in T2D subjects and 0.29 with non-diabetic controls, respectively.

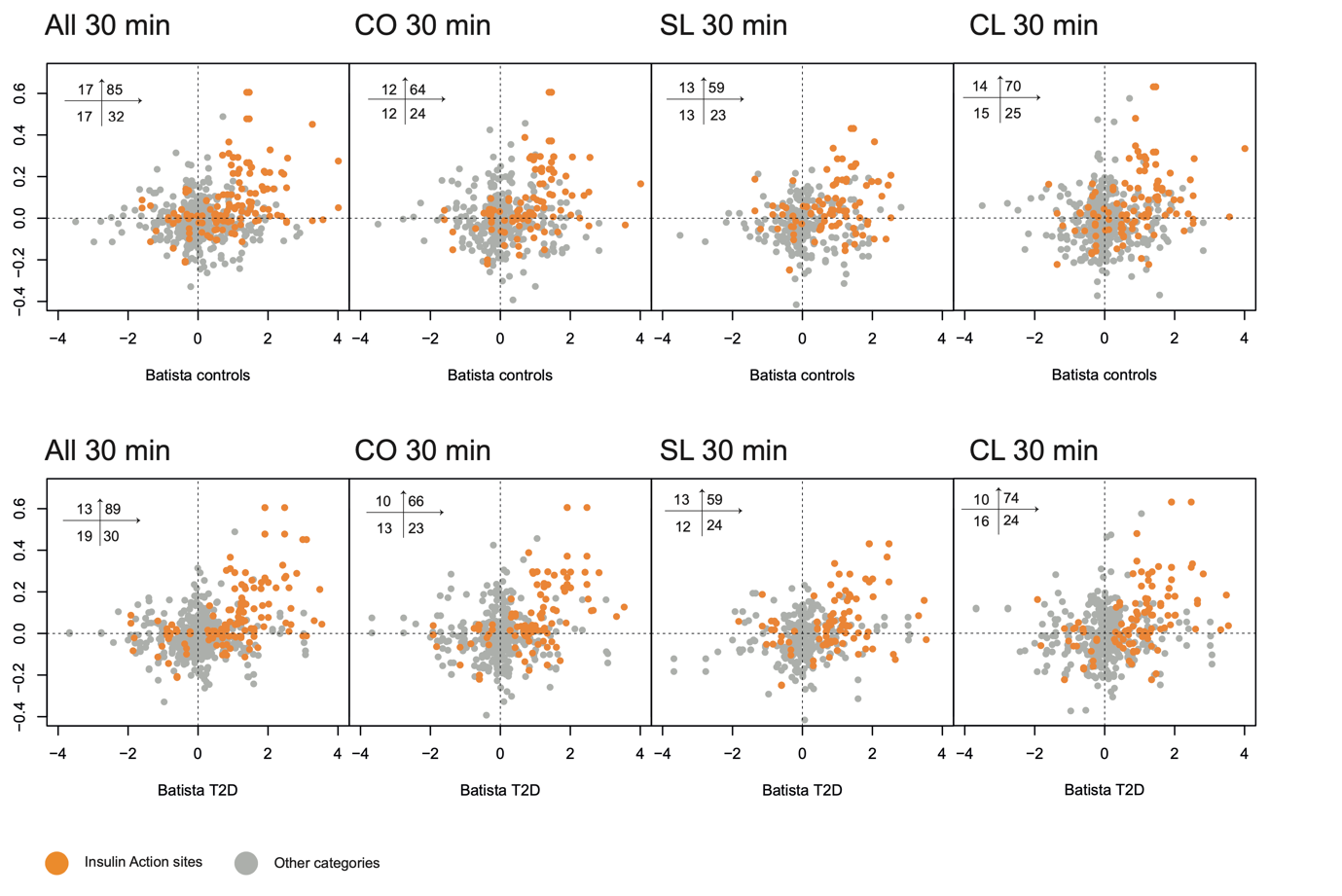

Scatter plots of log2 fold change of overlapped sites in our data on Y axis and Z-scored log2 intensity of overlapped sites in Batista *et al*. on X axis. The upper panel is comparison between our data and control subjects in Batista et al., whereas the lower panel is comparing to T2D subjects in Batista *et al*. Dots highlighted in orange are those identified as “Insulin Action” in Batista *et al*.

These comparable correlations were expected since the fold differences in the Z-scores were similar between T2D subjects and healthy subjects in the iMyos data (Figure 3 of the paper). Stratifying the comparison by our subject groups (CO/CL/SL), we did not observe particularly outstanding correlations in the insulin response for any of the three groups with the iMyos data. Limiting the comparison of matching pairs in the “insulin action” subset in Batista *et al*. (N=151), the Pearson correlation increased to 0.50 with T2D subjects and 0.42 with their controls. Within each group, we found the highest correlation with T2D and controls in CO (0.48 and 0.43), followed by CL (0.45 and 0.36) and SL (0.44 and 0.36), respectively.

**Detailed methods for the phosphoproteomics analysis of CDK1 inhibited myotubes**

The phosphoproteomics analysis of CDK1 inhibited myotubes derived from three biopsy donors was performed in an independent proteomics laboratory. Therefore there were minor differences in the sample preparation and LC-MS analysis steps, and . Here we provide the details here.

**Sample preparation**

Myotube cell pellets from three different patient-derived cultures with 12 conditions (4 treatment conditions (DMSO control, CDK1 inhibition by RO-3306 or knockdown using siNTC or siCDK1) and 3 time-points (0, 5, 30 min) per condition) were harvested. 200 µl of 8 M urea lysis buffer was added into each sample to denature proteins. Samples were sonicated for 3 cycles (15 s pulse and 45 s pause per cycle) to shear DNA and increase protein solubility. Total protein quantification was performed using Pierce BCA Protein Assay Kit (Thermo Fisher Scientific, Cat. No.: 23225) and 60 µg of protein was taken from each sample for proteomics experiment. Samples were reduced and alkylated with 20 mM TCEP and 55 mM CAA at room temperature in the dark, followed by digestion with 2 µg of endoproteinase LysC for 4 h and 2 µg trypsin overnight at 25 °C. Samples were checked for digestion efficiency before proceeding to the next step to ensure all peptides were digested effectively. After digestion, the samples were acidified and desalted with a 30 mg Oasis HLB plate (Waters, Cat. No.: WAT058951). A pooled sample was prepared by aliquoting equal amount of desalted peptides (2 µg) from all 48 samples. All 12 samples from the same patient culture and 4 replicates of the pooled sample were labelled together with TMTpro 16-plex for quantification. For each set of samples, sample complexity was reduced by step fractionation with 7 increasing acetonitrile concentrations of 14%, 18%, 21%, 24%, 27%, 32% and 60% in 10 mM ammonium formate, pH 10. After two repeats of acid wash (0.1% formic acid in water) to remove any remaining ammonium salts, the samples were loaded for mass spectrometry.

Remaining desalted samples from the preparation above (1.35 mg from each sample) were used for phospho-enrichment experiment. Samples were dried by vacuum centrifugation, resuspended in 6% trifluoroacetic acid (TFA) in 40% acetonitrile, and incubated with titanium dioxide (TiO_2_) beads (Titansphere, GL Sciences, Cat. No.: 5020-75000) for phosphoenrichment. This incubation was repeated once with fresh TiO_2_ beads by transferring the supernatant to new tubes. The beads were loaded over a C8 mesh, washed with 6% TFA in 10%, 40% and 60% acetonitrile sequentially, and phosphopeptides were eluted with 5% ammonia water followed by 5% ammonia water in 25% acetonitrile. Phosphopeptides were acidified and filtered to remove the TiO_2_ beads with a C8 frit prior to sample acquisition. Pooled samples and TMTpro 16-plex labelling were carried out as above mentioned. Each sample set was fractionated with 7 increasing acetonitrile concentration of 9%, 11.5%, 14%, 16.5%, 19%, 22% and 60% in 10 mM ammonium formate. After two repeats of acid wash (0.1% formic acid in water) to remove any remaining ammonium salts, the samples were loaded for mass spectrometry.

**Mass spectrometry**

Non-enriched total peptides were analyzed using reversed-phase liquid chromatography on the nLC1000 UHPLC system connected to Orbitrap Fusion Lumos mass spectrometer (Thermo Fisher Scientific). Each fraction was separated on 50 cm length x 75 µm internal diameter analytical C18 column (Thermo Scientific) in a 70 min gradient with a pre-programmed mixing of solvent A (0.1% formic acid in water) and solvent B (95% acetonitrile, 0.1% formic acid in water). Column was maintained at 50 °C and peptides were eluted using a 2–27% (v/v) acetonitrile gradient over 45 min, followed by an increase to 55% over the next 15 min, and to 95% over 5 min. The final mixture was maintained on the column for 5 min to elute all remaining peptides. Total run duration for each sample was 70 min at a constant flow rate of 300 nl/min.

Samples were ionized using 2.1 kV and 300 °C at the nanospray source. The following acquisition parameters were applied: data-dependent acquisition in positive mode with MS1 survey scan of 60,000 resolution over a scan range of 350–1550 *m/z*, automatic gain control (AGC) target: 8e5 ions, and maximum ion injection time: 50 ms. Precursors with charges 2–7 and having the highest ion counts in each MS1 scan were further fragmented using higher-energy collision-induced dissociation (HCD) at 42% normalized collision energy. MS2 was carried out using Orbitrap analyzer at 50,000 resolution, isolation window: 1 *m/z*, AGC target: 1e5 ions, and maximum IT: 80 ms. Precursors used for MS2 scans were excluded for 45 s to minimize re-sampling of high abundance peptides. The MS1–MS2 cycles were repeated every 3 s until completion of the run.

Phosphoenriched samples were subjected for mass spectrometry analysis using similar settings with Orbitrap Eclipse mass spectrometer (Thermo Fisher Scientific). Phosphopeptides were eluted using 2–30% (v/v) acetonitrile gradient over 70 min, followed by an increase to 45% over the next 10 min, and to 95% over 5 min. The final mixture was maintained on the column for 5 min to elute all remaining peptides. Total run duration was 90 min at a constant flow rate of 300 nl/min. The following MS parameters were changed: MS1 AGC target: 4e5, maximum IT: 100 ms; precursor was dynamically excluded for 60 s; MS2 isolation window: 1.2 *m/z*, AGC target: 7.5e4 ions, and maximum IT: 100 ms.

**Proteomics data analysis**

Proteins were identified using Proteome Discoverer™ (v3.0, Thermo Fisher Scientific). Raw mass spectra were searched against human primary protein sequences retrieved from UniProt (11 June 2019). Carbamidomethylation on Cys was set as a fixed modification; deamidation of Asn and Gln, acetylation on protein N termini, Met oxidation, and phosphorylation of Ser, Thr, and Tyr were set as dynamic modifications for the search. Trypsin/P was set as the digestion enzyme and was allowed up to three missed cleavage sites. Precursors and fragments were accepted if they had a mass error within 10 ppm and 0.06 Da, respectively. Peptides were matched to spectra at a false discovery rate (FDR) of 1% (strict) and 5% (relaxed) against the decoy database.

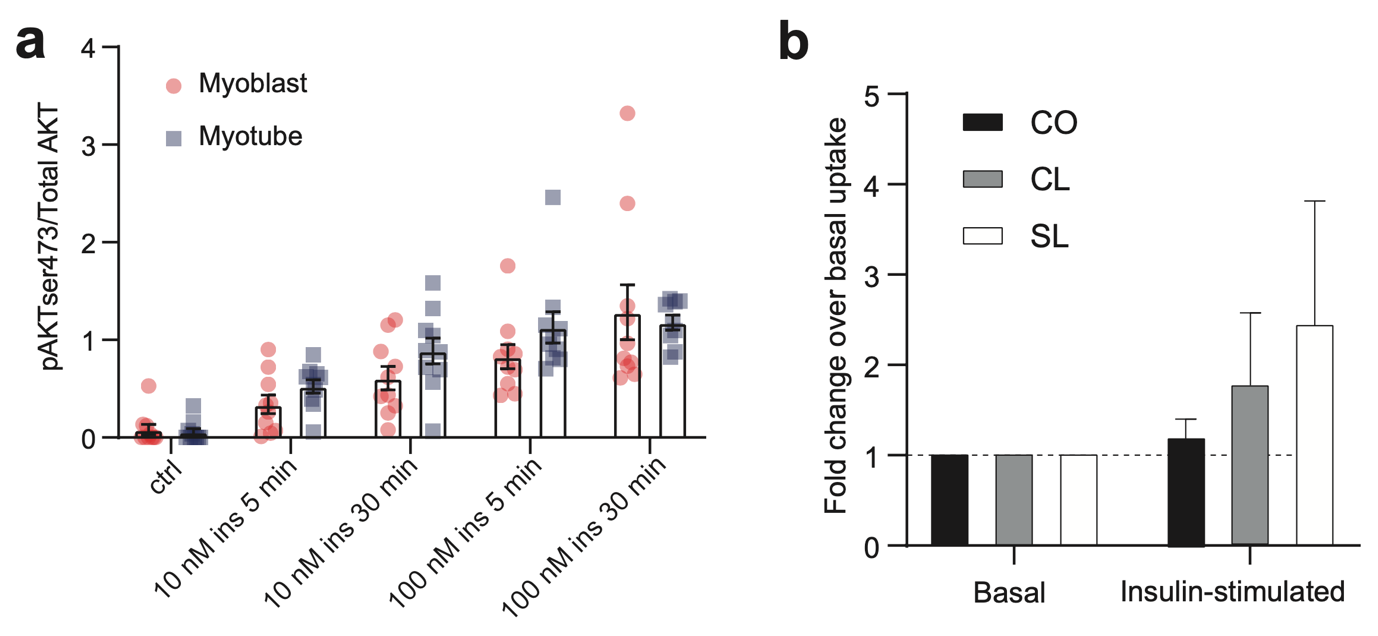

**Supplementary Figure 1. (a)** Densitometric analysis of p-Akt protein levels in myotubes isolated from subjects before (Ctrl) or after treatment (10 nM or 100 nM insulin) at 5 and 30 minutes. Graph displays mean p-Akt levels in arbitrary units (AU) normalized against total AKT protein levels. Error bars are expressed as mean ± SEM (n=9). **(b)** Insulin-stimulated glucose uptake (10 nM insulin) quantified by measuring luminescence signal proportional to concentration of 2DG taken up by myotubes. Data are presented as fold change over basal uptake (without insulin treatment) and expressed as mean ± SEM (n=3).

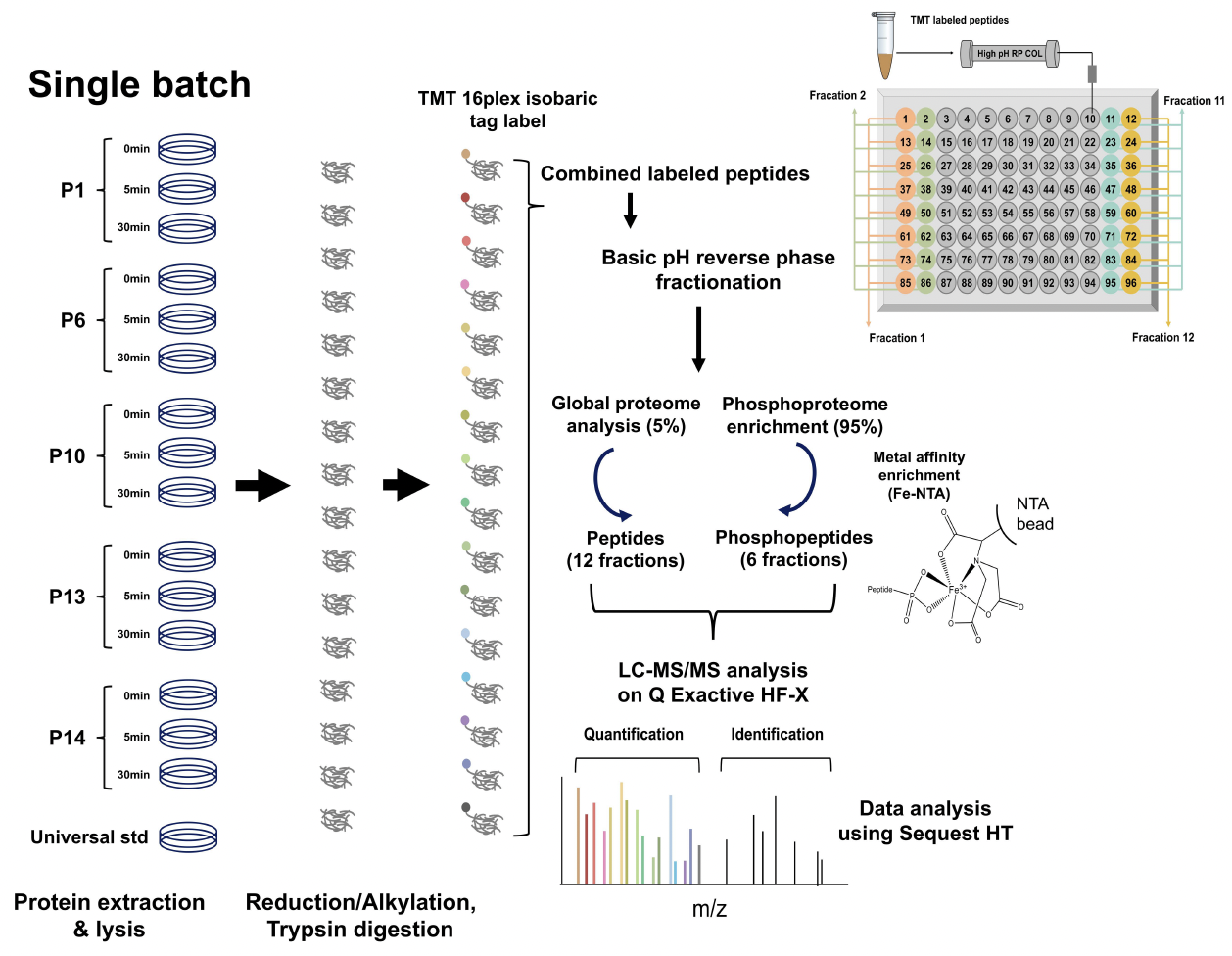

**Supplementary Figure 2**. Overall experimental design of mass spectrometric data acquisition for the global proteomics and phosphoproteomics analysis of the 90 myotube samples. The samples were allocated to 6 experimental batches (16-plex TMT labelling). The TMT-labelled samples were then combined and fractionated into twelve fractions for LC-MS/MS analysis. From the fractionated samples, 5% of the fractionated samples were used for the global proteome analysis and the other 95% of the fractionated samples were used for the phosphoproteome analysis following iron-chelate, ferric nitrilotriacetate (Fe-NTA) phosphopeptide enrichment.

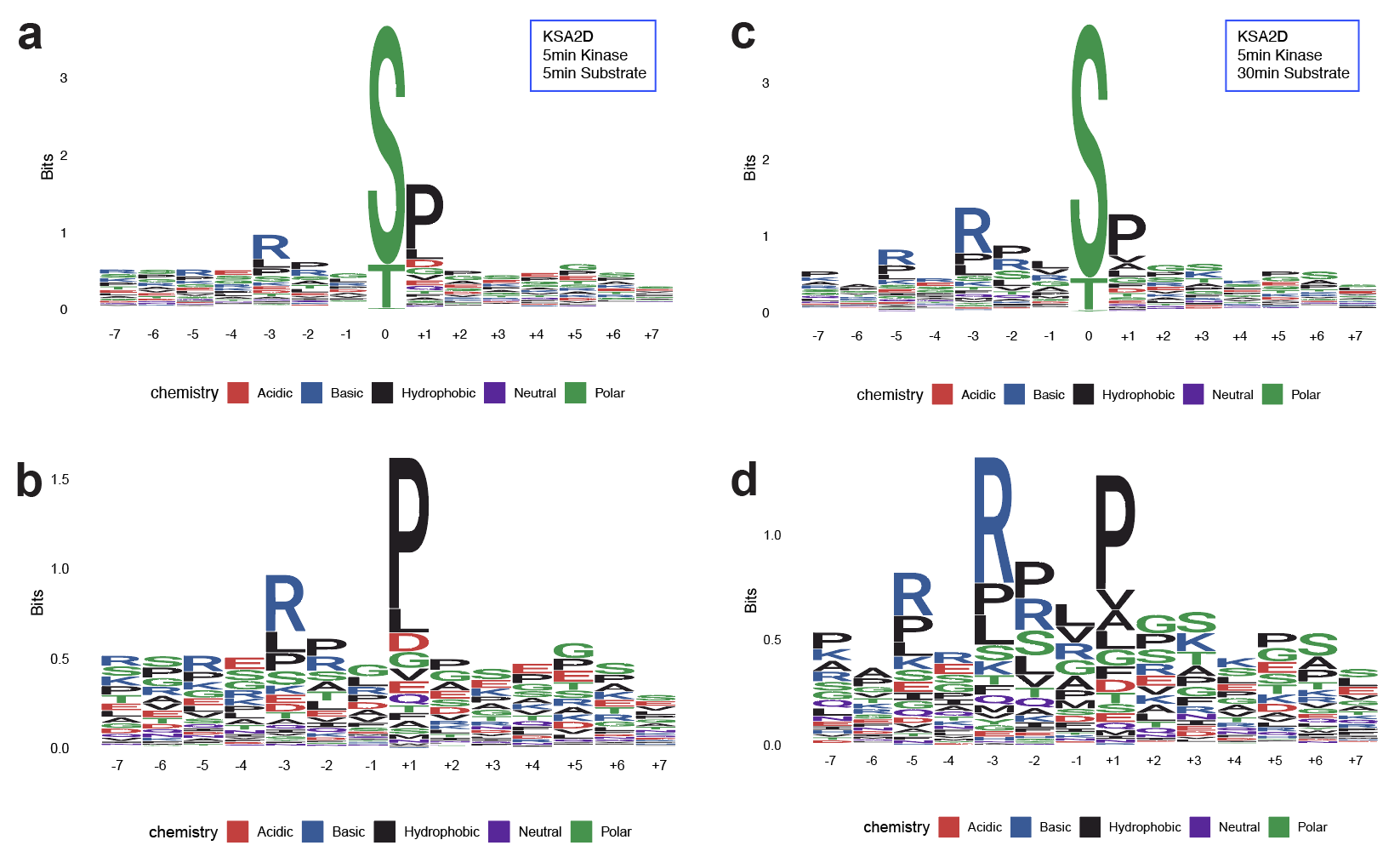

**Supplementary Figure 3.** Sequence motif logo plot from the motif enrichment analysis of the +/-7mer sequence windows around the phosphorylation sites identified by the KSA2D analysis for kinase abundance change at 5 min and substrate phosphorylation change at 5 min **(a)**, with the same logo plot rescaled after discarding the central S/T/Y **(b)**. The sequence motif logo plot from the enrichment analysis of the substrates identified by the KSA2D analysis of kinase abundance changes at 5 min and substrate phosphorylation changes at 30 min with and without the central S/T/Y (**c**, **d** respectively). The plots show specific enrichment of the minimal kinase motifs (S/T)-P and R-x-x-S/T.

­­
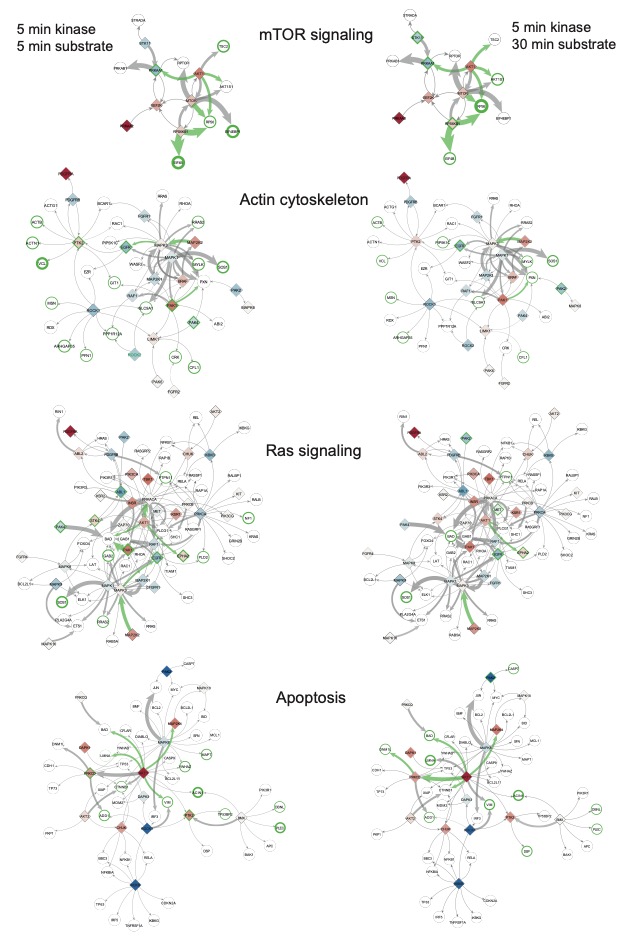

**Supplementary Figure 4.** Pathway-specific networks highlighting kinase-substrate pairs with significant positive co-variation in all thirty subjects (KSA 2D analysis results, at local false discovery rate 0.05) in green edges. Edge thickness is proportional to the number of possible phosphorylation sites seen in our data between the kinase and the substrate protein. Thin edges indicate theoretical phosphorylation relationships not recovered with our data. Node colors in diamonds show kinase protein abundance change. Green labels represent significant kinase abundance changes. Green node borders indicate occurrence of significant site level change of the corresponding substrate, and the border thickness is proportional to the number of significant sites.

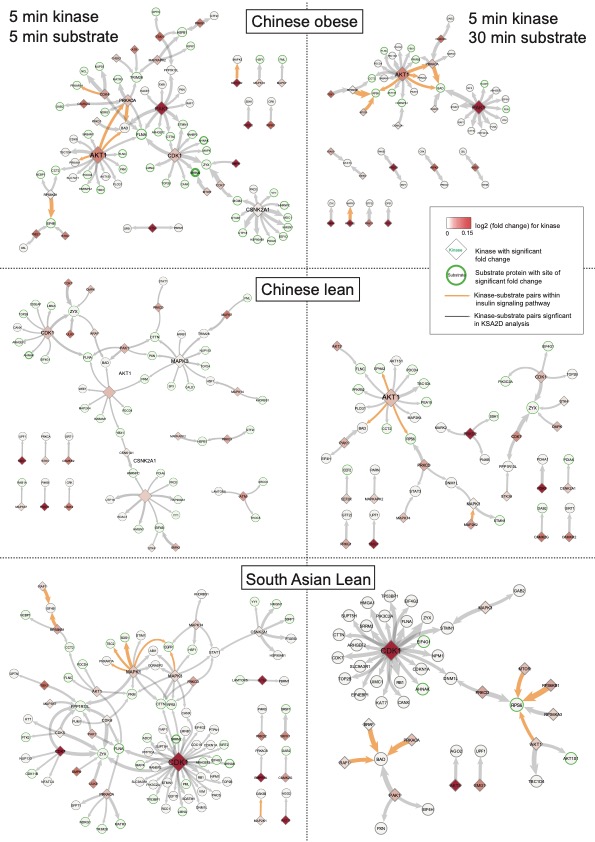

**Supplementary Figure 5.** Network of kinase-substrate pairs with significant positive co-variation in Chinese obese (CO), Chinese lean (CL) and South Asian lean (SL) subgroups. (2D analysis results, at local false discovery rate 0.05). Significant relationships of 5 min kinase with 5 min substrate in **(a)** and 5 min kinase with 30 min substrate in **(b).** Each edge represents phosphorylation of a substrate (sites omitted, in round circles) by a previously validated kinase (in diamonds). Edge thickness is proportional to the number of possible phosphorylation sites seen in our data between the kinase and the substrate protein. Node color in diamonds shows kinase protein abundance change. Green label represents significant kinase abundance change. Green node border indicates occurrence of significant site level change of the corresponding substrate, and the border thickness is proportional to the number of significant sites. Label and node size are proportional to the degree of connectivity in the network. Orange edge represents known kinase-substrate relationship in insulin signalling pathway.

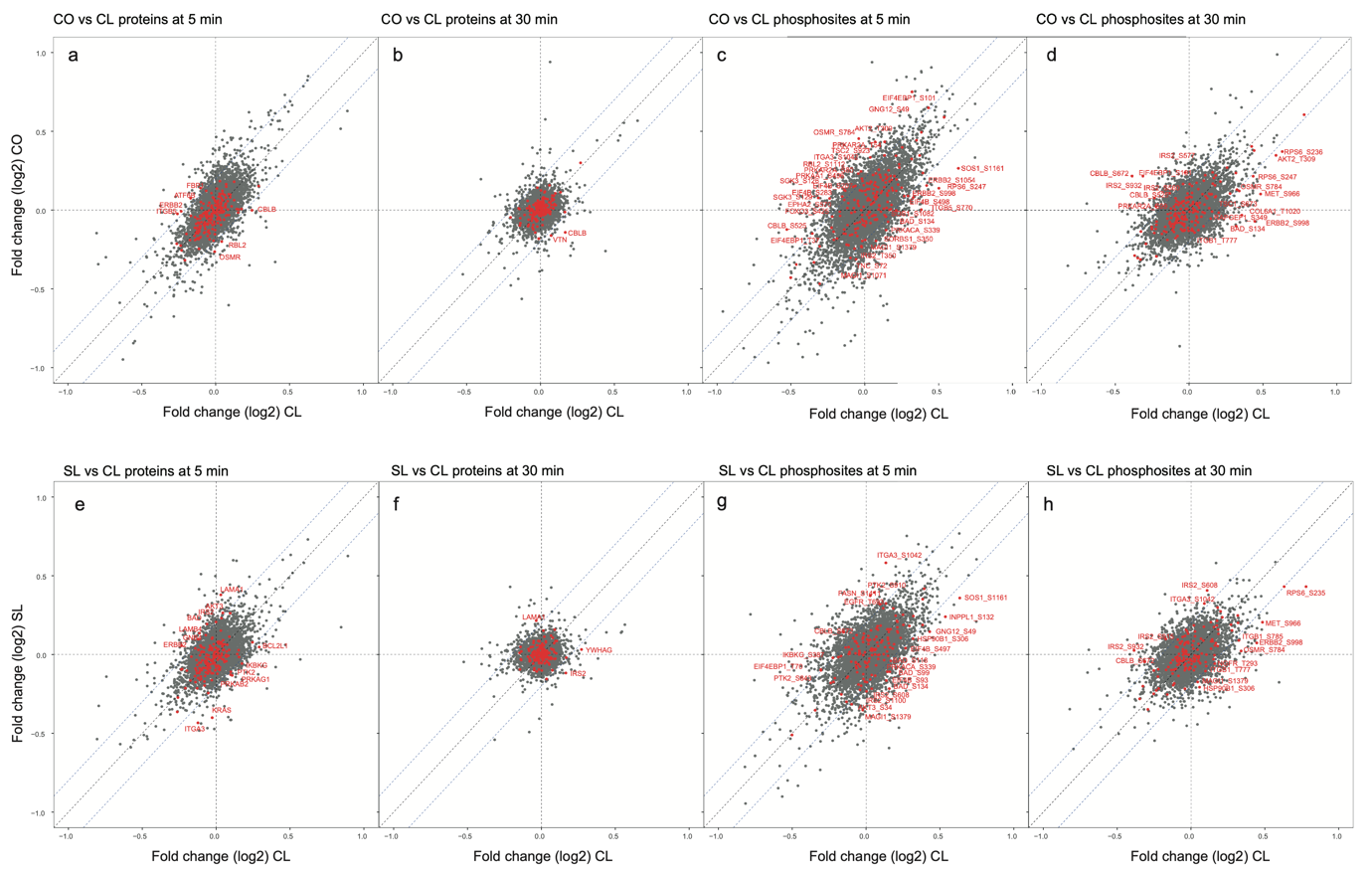
­

**Supplementary Figure 6**. Differences in the protein abundance and phosphorylation changes between groups. Log2 fold change (threshold 0.2 in absolute value) are plotted for CO and SL against CL at the two time points. Red dots are proteins or phosphorylation sites beyond the threshold (log2 fold change difference >0.2) in the insulin signalling pathway and the PI3K-AKT signalling pathway.
